# Transcripts with high distal heritability mediate genetic effects on complex metabolic traits

**DOI:** 10.1101/2024.09.26.613931

**Authors:** Anna L. Tyler, J. Matthew Mahoney, Mark P. Keller, Candice N. Baker, Margaret Gaca, Anuj Srivastava, Isabela Gerdes Gyuricza, Madeleine J. Braun, Nadia A. Rosenthal, Alan D. Attie, Gary A. Churchill, Gregory W. Carter

## Abstract

Although many genes are subject to local regulation, recent evidence suggests that complex distal regulation may be more important in mediating phenotypic variability. To assess the role of distal gene regulation in complex traits, we combined multi-tissue transcriptomes with physiological outcomes to model diet-induced obesity and metabolic disease in a population of Diversity Outbred mice. Using a novel high-dimensional mediation analysis, we identified a composite transcriptome signature that summarized genetic effects on gene expression and explained 30% of the variation across all metabolic traits. The signature was heritable, interpretable in biological terms, and predicted obesity status from gene expression in an independently derived mouse cohort and multiple human studies. Transcripts contributing most strongly to this composite mediator frequently had complex, distal regulation distributed throughout the genome. These results suggest that trait-relevant variation in transcription is largely distally regulated, but is nonetheless identifiable, interpretable, and translatable across species.

## Introduction

Evidence from genome-wide association studies (GWAS) suggests that most heritable variation in complex traits is mediated through regulation of gene expression. The majority of trait-associated variants lie in gene regulatory regions ^1–7^, suggesting a relatively simple causal model in which a variant alters the homeostatic expression level of a nearby gene which, in turn, alters a trait. Statistical methods such as transcriptome-wide association studies (TWAS) ^8–11^ and summary data-based Mendelian randomization (SMR) ^10^ have used this idea to identify genes associated with multiple disease traits ^12–15^. However, despite the great promise of these methods, explaining trait effects with local gene regulation has been more difficult than initially assumed ^16;17^. Although trait-associated variants typically lie in non-coding, regulatory regions, these variants often have no detectable effects on gene expression ^16^ and tend not to co-localize with expression quantitative trait loci (eQTLs) ^17;18^. These observations suggest that the relationship among genetic variants, gene expression, and organism-level traits is more complex than the simple, local model.

In recent years the conversation around the genetic architecture of common disease traits has been addressing this complexity, and there is increased interest in distal effects as potential drivers of trait variation ^18–20;15^. In particular, the omnigenic model posits that trait-driving genes are cumulatively influenced by many distal variants. In this view, the heritable transcriptomic signatures driving clinical traits are an emergent state arising from the myriad molecular interactions defining and constraining gene expression. Consistent with this view, it has been suggested that part of the difficulty in explaining trait variation through local eQTLs may arise in part because gene expression is not measured in the appropriate cell types ^16^, or cell states ^21^, and thus local eQTLs influencing traits cannot be detected in bulk tissue samples. This context dependence emphasizes the essential role of complex regulatory and tissue networks in mediating variant effects. The mechanistic dissection of complex traits in this model is more challenging because it requires addressing network-mediated effects that are weaker and greater in number. However, the comparative importance of distal effects over local effects is currently only conjectured and extremely challenging to address in human populations.

To assess the role of wide-spread distal gene regulation in the genetic architecture of complex traits, we used genetically diverse mice as a model system. In mice we can obtain simultaneous measurements of the genome, transcriptome, and phenome in all individuals. We used diet-induced obesity and metabolic disease as an archetypal example of a complex trait. In humans, these phenotypes are genetically complex with hundreds of variants mapped through GWAS ^22;23^ that are known to act through multiple tissues ^24;25^. Likewise in mice, metabolic traits are also genetically complex ^26^ and synteny analysis implicates a high degree of concordance in the genetic architecture between species ^26;12^. Furthermore, in contrast to humans, in mice we have access to multiple disease-relevant tissues in the same individuals with sufficient numbers for adequate statistical power.

We generated two complementary data sets: a discovery data set in a large population of Diversity Outbred (DO) mice ^27^, and an independent validation data set derived by crossing inbred strains from the Collaborative Cross (CC) recombinant inbred lines ^28^ to form CC recombinant inbred intercross (CC-RIX) mice. Both populations were maintained on a high-fat, high-sugar diet to model diet-induced obesity and metabolic disease ^12^.

The DO population and CC recombinant inbred lines were derived from the same eight inbred founder strains: five classical lab strains and three strains more recently derived from wild mice ^27^, representing three subspecies and capturing 90% of the known variation in laboratory mice ^29^. The DO mice are maintained with a breeding scheme that ensures equal contributions from each founder across the genome thus rendering almost the whole genome visible to genetic inquiry and maximizing power to detect eQTLs^27^. The CC mice were initially intercrossed to recombine the genomes from all eight founders, and then inbred for at least 20 generations to create recombinant inbred lines ^28;30;29^. Because these two populations have common ancestral haplotypes but highly distinct kinship structure, we could directly and unambiguously compare the local genetic effects on gene expression at the whole-transcriptome level while varying the population structure driving distal regulation.

In the DO population, we paired clinically relevant metabolic traits, including body weight and plasma levels of insulin, glucose and lipids ^12^, with transcriptome-wide gene expression in four tissues related to metabolic disease: adipose tissue, pancreatic islets, liver, and skeletal muscle. We measured similar metabolic traits in a CC-RIX population and gene expression from three of the four tissues used in the DO: adipose tissue, liver, and skeletal muscle. Measuring gene expression in multiple tissues is critical to adequately assess the extent to which local gene regulation varies across the tissues and whether such variability might account for previous failed attempts to identify trait-relevant local eQTLs. The CC-RIX carry the same founder alleles as the DO. Thus, local gene regulation is expected to match between the populations. However, because the alleles are recombined throughout the genome, distal effects are expected to vary from those in the DO, allowing us to directly assess the role of distal gene regulation in driving trait-associated transcript variation. To mechanistically dissect distal effects on metabolic disease, we developed a novel dimension reduction framework called high-dimensional mediation analysis (HDMA) to identify the heritable transcriptomic signatures driving trait variation, which we compared between mouse populations and to human data sets with measured adipose gene expression. Together, these data enable a comprehensive view into the genetic architecture of metabolic disease.

## Results

### Genetic variation contributed to wide phenotypic variation

Although the environment was consistent across the DO mice, the genetic diversity present in this population resulted in widely varying distributions across physiological measurements (Fig. 1). For example, body weights of adult individuals varied from less than the average adult C57BL/6J (B6) body weight to several times the body weight of a B6 adult in both sexes (Males: 18.5 - 69.1g, Females: 16.0 - 54.8g) (Fig. 1A). Fasting blood glucose (FBG) also varied considerably (Fig. 1B), although few of the animals had FBG levels that would indicate pre-diabetes (19 animals, 3.8%), or diabetes (7 animals, 1.4%) according to previously developed cutoffs (pre-diabetes: FBG *≥* 250 mg/dL, diabetes: FBG *≥* 300, mg/dL) ^31^. Males had higher FBG than females on average (Fig. 1C) as has been observed before suggesting either that males were more susceptible to metabolic disease on the high-fat, high-sugar (HFHS) diet, or that males and females may require different thresholds for pre-diabetes and diabetes.

**Figure 1:**
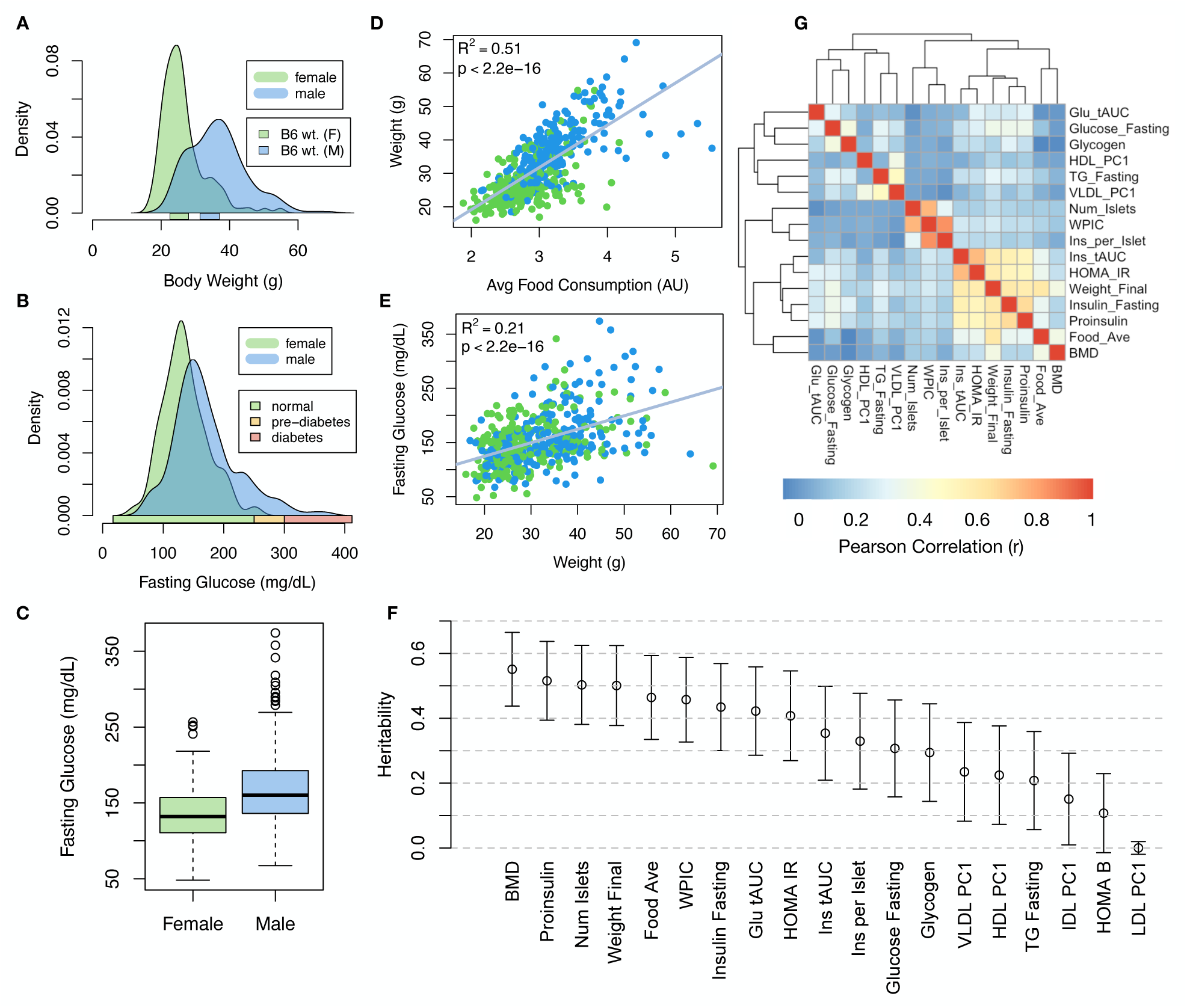
Clinical overview. **A.** Distributions of body weight in the diversity outbred mice. Sex is indicated by color. The average B6 male and female adult weights at 24 weeks of age are indicated by blue and green bars on the x-axis. **B.** The distribution of fasting glucose across the population split by sex. Normal, pre-diabetic, and diabetic fasting glucose levels for mice are shown by colored bars along the x-axis. **C.** Males had higher fasting blood glucose on average than females. **D.** The relationship between food consumption and body weight for both sexes. **E.** Relationship between body weight and fasting glucose for both sexes. **F.** Heritability estimates for each physiological trait. Bars show standard error of the estimate. **G.** Correlation structure between pairs of physiological traits. BMD - bone mineral density, WPIC - whole pancreas insulin content, Glu tAUC - glucose total area under the curve, HOMA IR - homeostatic measurement of insulin resistance, HOMA B - homeostatic measure of beta cell health, VLDL - very low-density lipoprotein, LDL - low-density lipoprotein, IDL - intermediate density lipoprotein, HDL - high-density lipoprotein, TG - triglyceride.

Body weight was strongly positively correlated with food consumption (Fig. 1D *R*^2^ = 0.51, *p <* 2.2 *×* 10*^−^*^16^) and FBG (Fig. 1E, *R*^2^ = 0.21, *p <* 2.2 *×* 10*^−^*^16^) suggesting a link between behavioral factors and metabolic disease. However, the heritability of this trait and others (Fig. 1F) indicates that genetics contribute substantially to correlates of metabolic disease in this population.

The trait correlations (Fig. 1G) showed that most of the metabolic trait pairs were only modestly correlated, which, in conjunction with the trait decomposition (Supp. Fig. S1), suggests complex relationships among the measured traits and a broad sampling of multiple heritable aspects of metabolic disease including overall body weight, glucose homeostasis, and pancreatic function.

### Distal Heritability Correlated with Phenotype Relevance

To comprehensively assess the genetic control of gene expression in metabolic disease we measured overall gene expression via bulk RNA-Seq in adipose, islet, liver, and skeletal muscle in the DO cohort (Supp. Fig. S2). We performed eQTL analysis using R/qtl2^32^ (Methods) and identified both local and distal eQTLs for transcripts in each of the four tissues (Supp. Fig. S2B-E). Significant local eQTLs far outnumbered distal eQTLs (Supp. Fig. S2F) and tended to be shared across tissues (Supp. Fig. S2G) whereas the few significant distal eQTLs we identified tended to be tissue-specific (Supp. Fig. S2H)

We calculated the heritability of each transcript in terms of local and all non-local (distal) genetic factors (Methods). Overall, local and distal genetic factors contributed approximately equally to transcript abundance. In all tissues, both local and distal factors explained between 8 and 18% of the variance in the median transcript (Fig, 2A).

To assess the importance of genetic regulation of transcript levels to clinical traits, we compared the local and distal heritabilities of transcripts to their trait relevance, defined as the maximum trait correlation for each transcript. The local heritability of transcripts was negatively correlated with their trait relevance (Fig. 2B), suggesting that the more local genotype influenced transcript abundance, the less effect this variation had on the measured traits. Conversely, the distal heritability of transcripts was positively correlated with trait relevance (Fig. 2C). That is, transcripts that were more highly correlated with the measured traits tended to be distally, rather than locally, heritable. Importantly, this pattern was consistent across all tissues. This finding is consistent with previous observations that transcripts with low local heritability explain more expression-mediated disease heritability than transcripts with high local heritability ^19^. However, the positive

**Figure 2:**
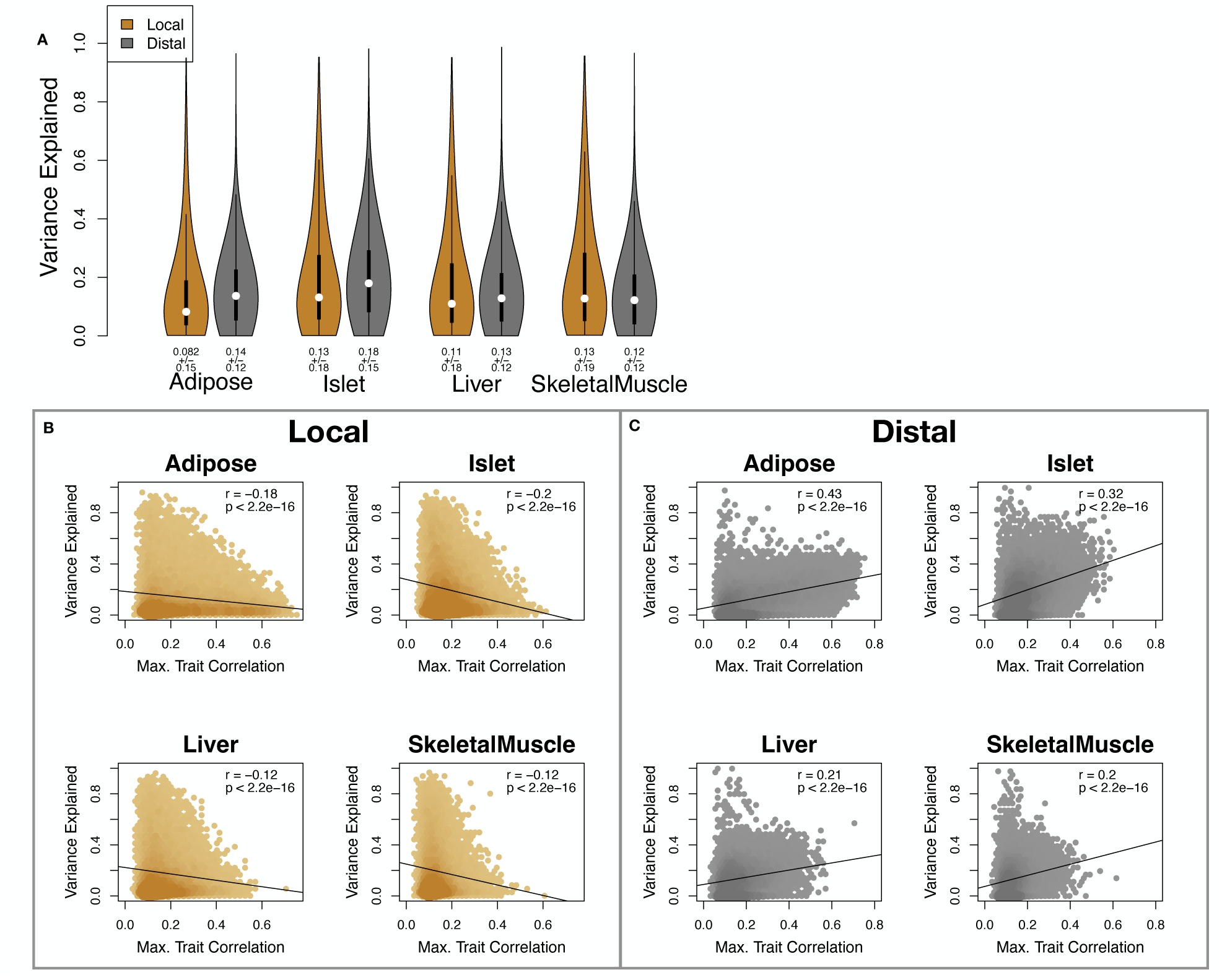
Transcript heritability and trait relevance. **A.** Distributions of distal and local heritability of transcripts across the four tissues. Overall local and distal factors contribute equally to transcript heritability. The relationship between (**B.**) local and (**C.**) distal heritability and trait relevance across all four tissues. Here trait relevance is defined as the maximum correlation between the transcript and all traits. Local heritability was negatively correlated with trait relevance, and distal heritability is positively correlated with trait relevance. Pearson (*r*) and *p* values for each correlation are shown in the upper-right of each panel. relationship between trait correlation and distal heritability demonstrated further that there are diffuse genetic effects throughout the genome converging on trait-related transcripts.

### High-Dimensional Mediation Analysis identified a high-heritability composite trait that was mediated by a composite transcript

The above univariate analyses establish the importance of distal heritability for trait-relevant transcripts. However, the number of transcripts dramatically exceeds the number of phenotypes. Thus, we expect the heritable, trait-relevant transcripts to be highly correlated and organized according to coherent, biological processes representing the mediating endophenotypes driving clinical trait variation. To identify these endophenotypes in a theoretically principled way, we developed a novel dimension-reduction technique, high-dimension mediation analysis (HDMA), that uses the theory of causal graphical models to identify a transcriptomic signature that is simultaneously 1) highly heritable, 2) strongly correlated to the measured phenotypes, and 3) conforms to the causal mediation hypothesis (Fig. 3). HDMA projects the high-dimensional genome, transcriptome, and phenome data onto one-dimensional scores–a composite genome score (*G_C_*), a composite transcriptome score (*T_C_*), and a composite phenome score (*P_C_*)–and uses the univariate theory of mediation to constrain these projections to satisfy the hypotheses of perfect mediation, namely that upon controlling for the transcriptomic score, the genome score is uncorrelated to the phenome score. A complete mathematical derivation and implementation details for HDMA are available in Supp. Methods.

**Figure 3:**
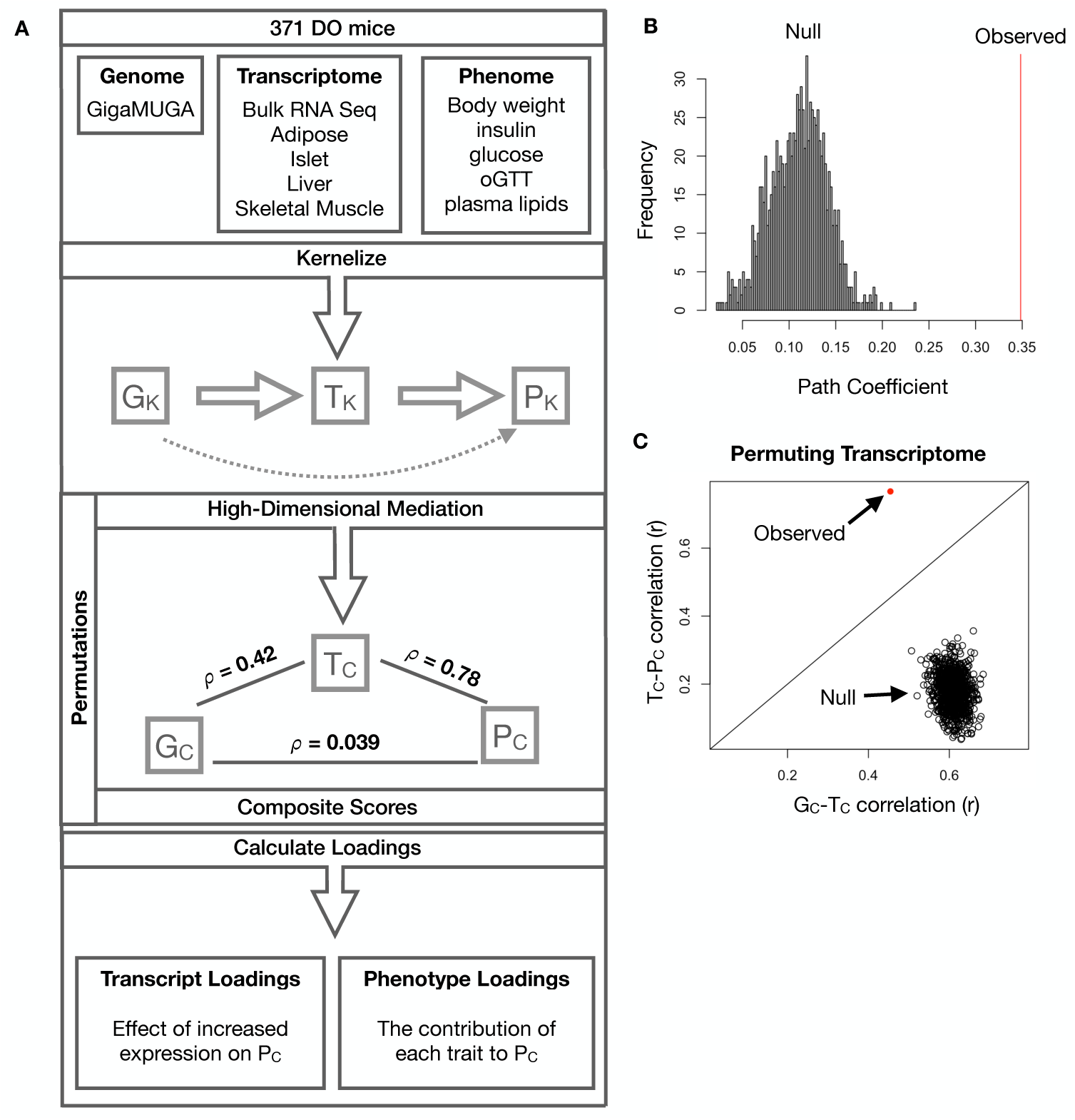
High-dimensional mediation. **A.** Workflow indicating major steps of high-dimensional mediation. The genotype, transcriptome, and phenotype matrices were independently normalized and converted to kernel matrices representing the pairwise relationships between individuals for each data modality (*K_G_* = genome kernel, *K_T_*= transcriptome kernel; *K_P_*= phenome kernel). High-dimensional mediation was applied to these matrices to maximize the direct path *G → T → P* , the mediating pathway (arrows), while simultaneously minimizing the direct *G → P* pathway (dotted line). The composite vectors that resulted from high-dimensional mediation were *G_c_*, *T_C_*, and *P_C_*. The partial correlations *ρ* between these vectors indicated perfect mediation. Transcript and trait loadings were calculated as described in the methods. **B.** The null distribution of the path coefficient derived from 10,000 permutations compared to the observed path coefficient (red line). **C.** The null distribution of the *G_C_*-*T_C_* correlation vs. the *T_C_*-*P_C_* correlation compared with the observed value (red dot). genetically determined variation in phenotype that is mediated through genetically determined variation in transcription.

Using HDMA we identifed the major axis of variation in the transcriptome that was consistent with mediating the effects of the genome on metabolic traits (Fig 3). Fig. 3A shows the partial correlations (*ρ*) between the pairs of these composite vectors. The partial correlation between *G_C_*and *T_C_*was 0.42, and the partial correlation between *T_C_* and *P_C_* was 0.78. However, when the transcriptome was taken into account, the partial correlation between *G_C_* and *P_C_* was effectively zero (0.039). *P_C_* captured 30% of the overall trait variance, and its estimated heritability was 0.71 *±* 0.084, which was higher than any of the measured traits (Fig. 1F). Thus, HDMA identified a maximally heritable metabolic composite trait and a highly heritable component of the transcriptome that are correlated as expected in the perfect mediation model.

As discussed in Supp. Methods, HDMA is related to a generalized form of CCA. Standard CCA is prone to over-fitting because in any two large matrices it can be trivial to identify highly correlated composite vectors ^33^. To assess whether our implementation of HDMA was similarly prone to over-fitting in a high-dimensional space, we performed permutation testing. We permuted the individual labels on the transcriptome matri× 10,000 times and recalculated the path coefficient, which is the correlation of *G_C_* and *T_C_* multiplied by the correlation of *T_C_* and *P_C_*. This represents the strength of the path from *G_C_* to *P_C_* that is putatively mediated through *T_C_*. The null distribution of the path coefficient is shown in Fig. 3B, and the observed path coefficient from the original data is indicated by a red line. The observed path coefficient was well outside the null distribution generated by permutations (*p <* 10*^−^*^16^). Fig. 3C illustrates this observation in more detail. Although we identified high correlations between *G_C_* and *T_C_*, and modest correlations between *T_C_* and *P_C_* in the null data (Fig 3C), these two values could not be maximized simultaneously in the null data. In contrast, the red dot shows that in the real data both the *G_C_*-*T_C_* correlation and the *T_C_*-*P_C_* correlation could be maximized simultaneously suggesting that the path from genotype to phenotype through transcriptome is highly non-trivial and identifiable in this case. These results suggest that these composite vectors represent

### Body weight and insulin resistance were highly represented in the expression-mediated composite trait

Each composite score is a weighted combination of the measured variables. The magnitude and sign of the weights, called loadings, correspond to the relative importance and directionality of each variable in the composite score. The loadings of each measured trait onto *P_C_*indicate how much each contributed to the composite phenotype. Body weight contributed the most (Fig. 4), followed by homeostatic insulin resistance (HOMA_IR) and fasting plasma insulin levels (Insulin_Fasting). We can thus interpret *P_C_* as an index of metabolic disease (Fig. 4B). Individuals with high values of *P_C_*have a higher metabolic disease index (MDI) and greater metabolic disease, including higher body weight and higher insulin resistance. We refer to *P_C_* as the MDI going forward. Traits contributing the least to the MDI were measures of cholesterol and pancreas composition. Thus, when we interpret the transcriptomic signature identified by HDMA, we are explaining primarily the putative transcriptional mediation of body weight and insulin resistance, as opposed to cholesterol measurements.

**Figure 4:**
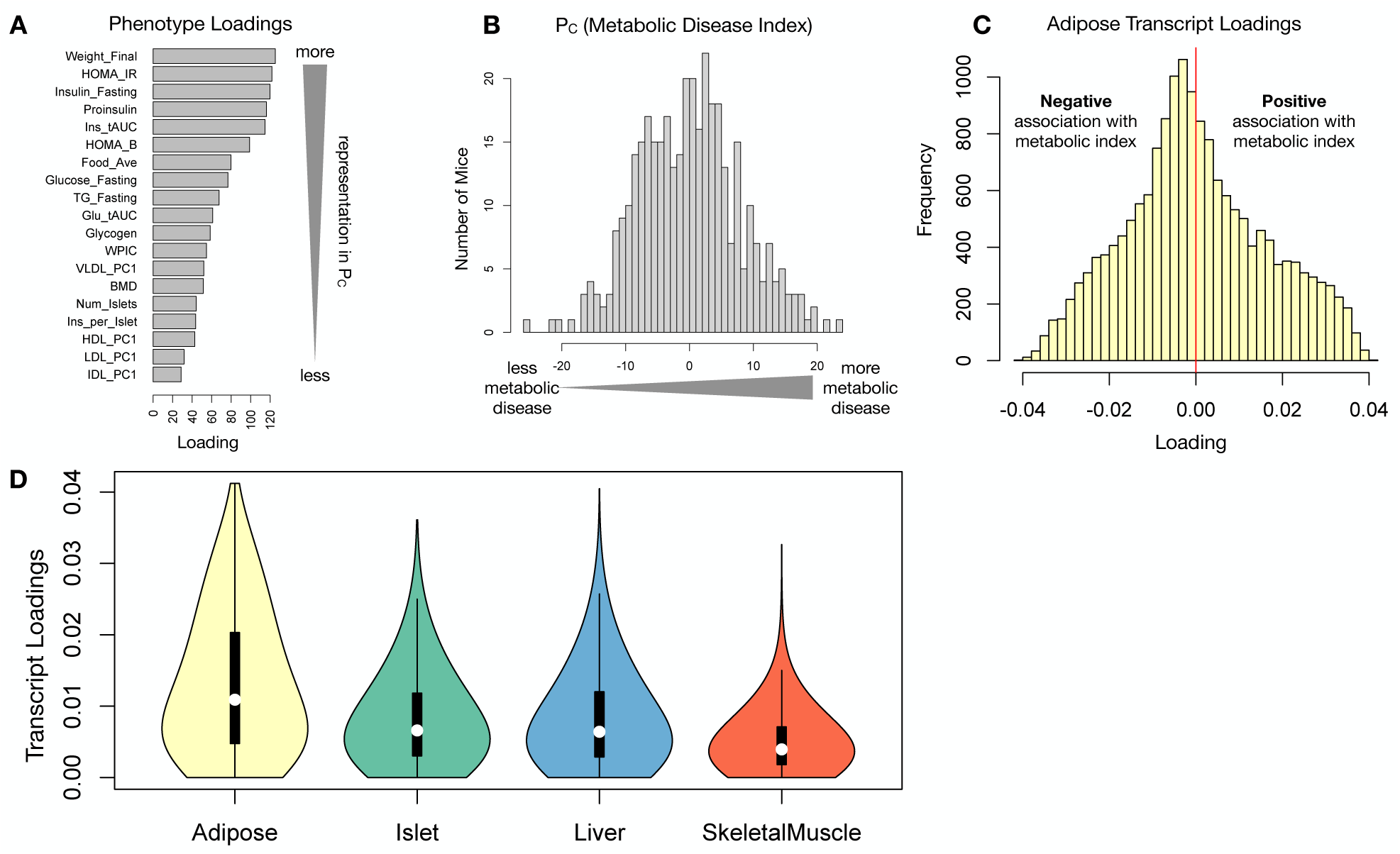
Interpretation of loadings. **A.** Loadings across traits. Body weight and insulin resistance contributed the most to the composite trait. **B.** Phenotype scores across individuals. Individuals with large positive phenotype scores had higher body weight and insulin resistance than average. Individuals with large negative phenotype scores had lower body weight and insulin resistance than average. **C.** Distribution of transcript loadings in adipose tissue. For transcripts with large positive loadings, higher expression was associated with higher phenotype scores. For transcripts with large negative loadings, higher expression was associated with lower phenotype scores. **D.** Distribution of absolute value of transcript loadings across tissues. Transcripts in adipose tissue had the largest loadings indicating that adipose tissue gene expression was a strong mediator of genotype on body weight and insulin resistance.

### High-loading transcripts had low local heritability, high distal heritability, and were linked mechanistically to obesity

We interpreted large loadings onto transcripts as indicating strong mediation of the effect of genetics on the MDI. Large positive loadings indicate that higher expression was associated with a higher MDI (i.e. higher risk of obesity and metabolic disease on the HFHS diet) (Fig. 4C). Conversely, large negative loadings indicate that high expression of these transcripts was associated with a lower MDI (i.e. lower risk of obesity and metabolic disease on the HFHS diet) (Fig. 4C). We used gene set enrichment analysis (GSEA) ^34;35^ to look for biological processes and pathways that were enriched at the top and bottom of this list (Methods).

In adipose tissue, both GO processes and KEGG pathway enrichments pointed to an axis of inflammation and metabolism (Figs. S3 and S4). GO terms and KEGG pathways associated with inflammation were positively associated with the MDI, indicating that increased expression in inflammatory pathways was associated with a higher burden of disease. It is well established that adipose tissue in obese individuals is inflamed and infiltrated by macrophages^36–40^, and the results here suggest that this may be a dominant heritable component of metabolic disease.

The strongest negative enrichments in adipose tissue were related to mitochondial activity in general, and thermogenesis in particular (Figs. S3 and S3). Genes in the KEGG oxidative phosphorylation pathway were almost universally negatively loaded in adipose tissue, suggesting that increased expression of these genes was associated with reduced MDI (Supp. Fig. S5). Consistent with this observation, it has been shown previously that mouse strains with greater thermogenic potential are also less susceptible to obesity on an obesigenic diet ^41^.

Transcripts associated with the citric acid cycle as well as the catabolism of the branched-chain amino acids (valine, leuceine, and isoleucine) were strongly enriched with negative loadings in adipose tissue (Supp. Figs. S3, S6 and S7). Expression of genes in both pathways (for which there is some overlap) has been previously associated with insulin sensitivity ^12;42;43^, suggesting that heritable variation in regulation of these pathways may influence risk of insulin resistance.

Looking a the 10 most positively and negatively loaded transcripts from each tissue, it is apparent that transcripts in the adipose tissue had the largest loadings, both positive and negative (Fig. 5A bar plot). This suggests that much of the effect of genetics on body weight and insulin reisistance is mediated through gene expression in adipose tissue. The strongest loadings in liver and pancreas were comparable, and those in skeletal muscle were the weakest (Fig. 5A), suggesting that less of the genetic effects were mediated through transcription in skeletal muscle. As expected, heritability analysis showed that transcripts with the largest loadings had higher distal heritability than local heritability (Fig. 5A heat map and box plot). This pattern contrasts with transcripts nominated by TWAS (Fig. 5B), which tended to have lower loadings, higher local heritability and lower distal heritability. Transcripts with the highest local heritability in each tissue (Fig. 5C) had the lowest loadings, consistent with our findings above (Fig. 2B).

**Figure 5:**
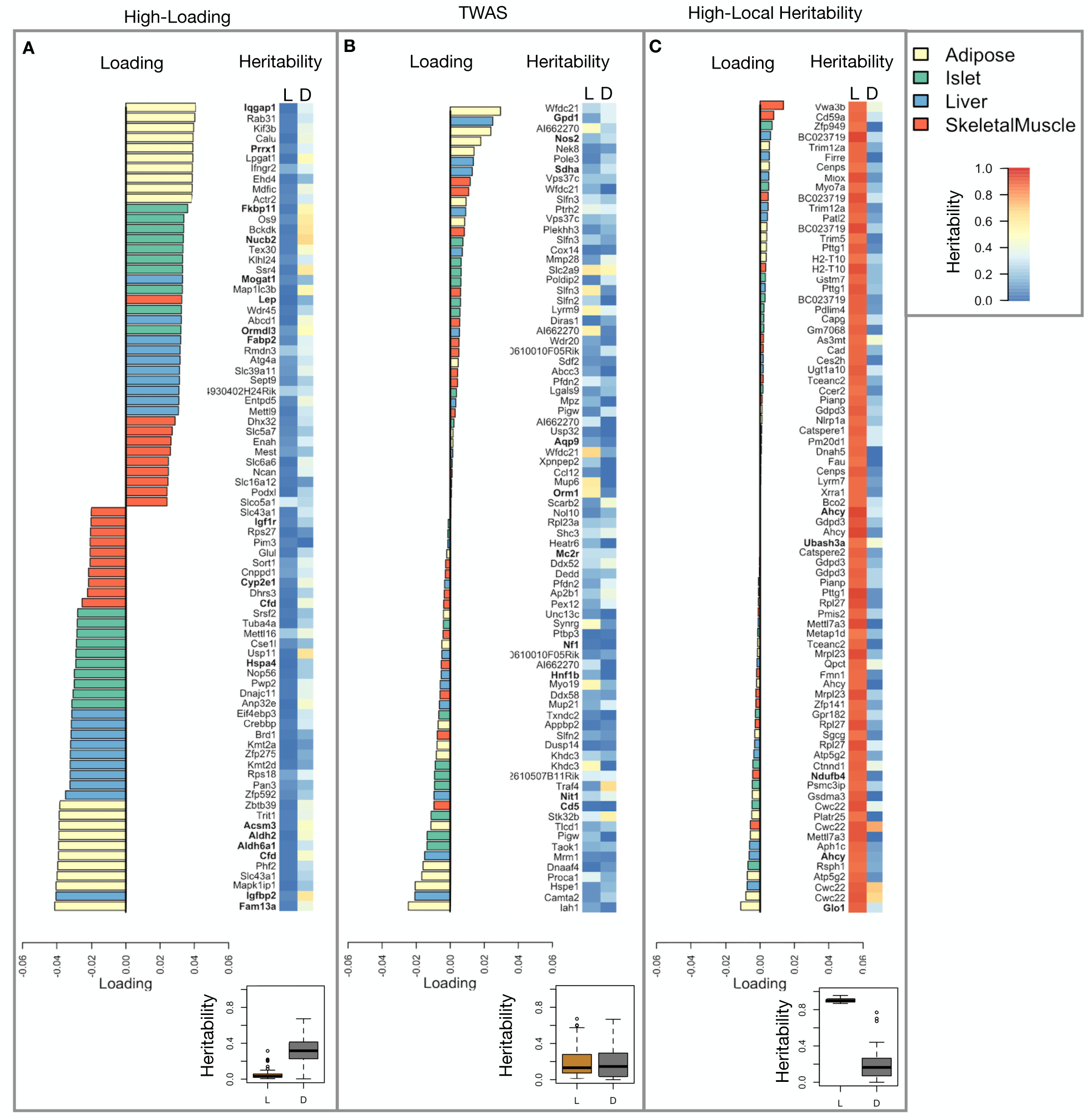
Transcripts with high loadings have high distal heritability and literature support. Each panel has a bar plot showing the loadings of transcripts selected by different criteria. Bar color indicates the tissue of origin. The heat map shows the local (L - left) and distal (D - right) heritability of each transcript. **A.** Loadings for the 10 transcripts with the largest positive loadings and the 10 transcripts with the largest negative loadings for each tissue. **B.** Loadings of TWAS candidates with the 10 largest positive correlations with traits and the largest negative correlations with traits across all four tissues. **C.** The transcripts with the largest local heritability (top 20) across all four tissues.

We performed a literature search for the genes in each of these groups along with the terms “diabetes”, “obesity”, and the name of the expressing tissue to determine whether any of these genes had previous associations with metabolic disease in the literature (Methods). Multiple genes in each group had been previously associated with obesity and diabetes (Fig. 5 bolded gene names). Genes with high loadings were most highly enriched for previous literature support. They were 2.4 times more likely than TWAS hits and 3.8 times more likely than genes with high local heritability to be previously associated with obesity or diabetes.

### Tissue-specific transriptional programs were associated with metabolic traits

Clustering of transcripts with top loadings in each tissue showed tissue-specific functional modules associated with obesity and insulin resistance (Fig. 6A) (Methods). The clustering highlights the importance of immune activation particularly in adipose tissue. The “mitosis” cluster had large positive loadings in three of the four tissues potentially suggesting system-wide proliferation of immune cells. Otherwise, all clusters were strongly loaded in only one or two tissues. For example, the lipid metabolism cluster was loaded most heavily in liver. The positive loadings suggest that high expression of these genes, particularly in the liver, was associated with increased metabolic disease. This cluster included the gene *Pparg*, whose primary role is in the adipose tissue where it is considered a master regulator of adipogenesis ^44^. Agonists of *Pparg*, such as thiazolidinediones, are FDA-approved to treat type II diabetes, and reduce inflammation and adipose hyptertrophy ^44^. Consistent with this role, the loading for *Pparg* in adipose tissue was negative, suggesting that higher expression was associated with leaner mice (Fig. 6B). In contrast, *Pparg* had a large positive loading in liver, where it is known to play a role in the development of hepatic steatosis, or fatty liver. Mice that lack *Pparg* specifically in the liver, are protected from developing steatosis and show reduced expression of lipogenic genes ^45;46^. Overexpression of *Pparg* in the livers of mice with a *Ppara* knockout, causes upregulation of genes involved in adipogenesis ^47^. In the livers of both mice and humans high *Pparg* expression is associated with hepatocytes that accumulate large lipid droplets and have gene expression profiles similar to that of adipocytes ^48;49^. The local and distal heritability of *Pparg* is low in adipose tissue suggesting its expression in this tissue is highly constrained in the population (Fig. 6B). However, the distal heritability of *Pparg* in liver is relatively high suggesting it is complexly regulated and has sufficient variation in this population to drive variation in phenotype. Both local and distal heribatility of *Pparg* in the islet are relatively high, but the loading is low, suggesting that variability of expression in the islet does not drive variation in MDI. These results highlight the importance of tissue context when investigating the role of heritable transcript variability in driving phenotype. Gene lists for all clusters are available in Supp. File 1.

**Figure 6:**
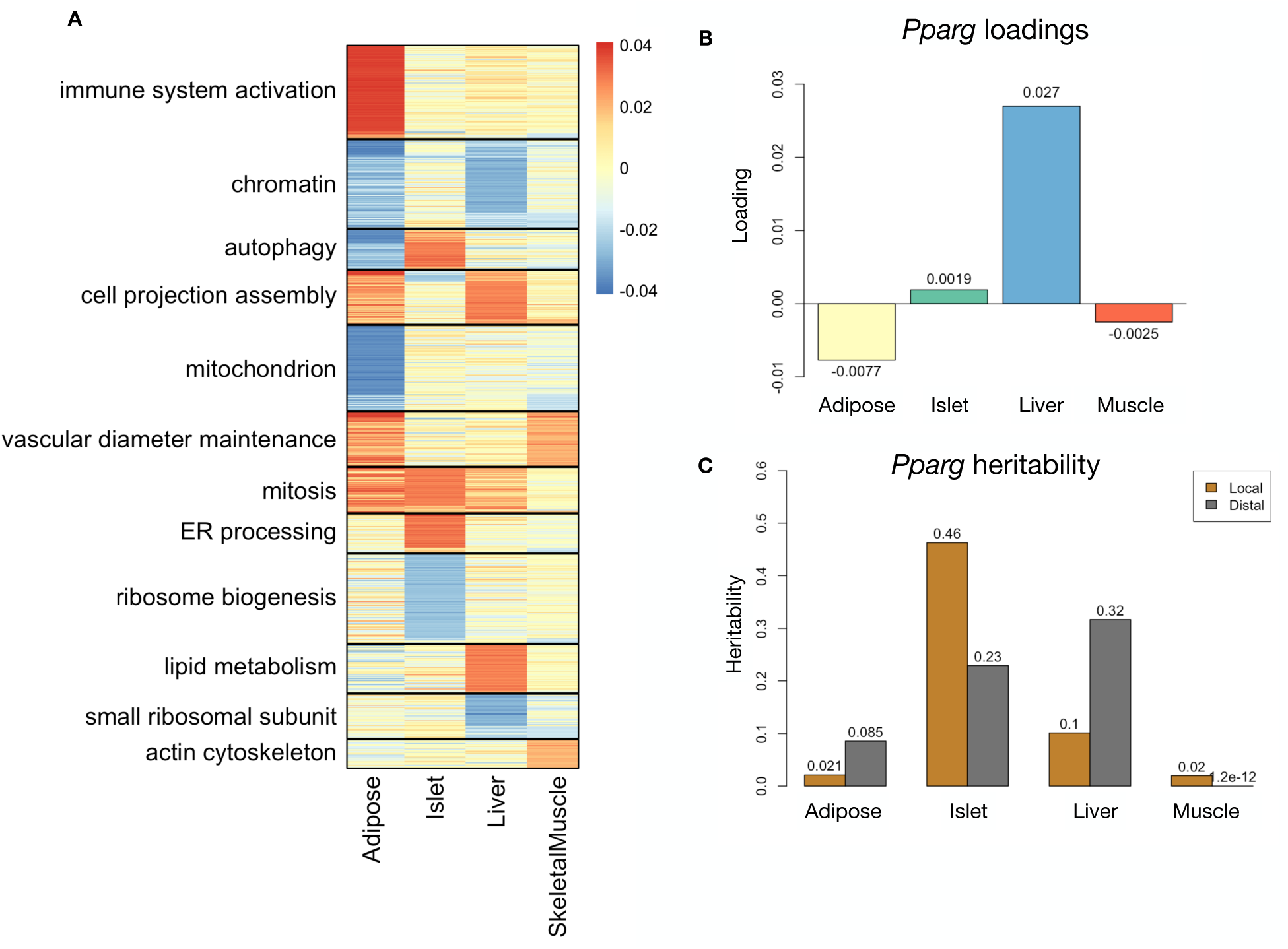
Tissue-specific transcriptional programs were associated with obesity and insulin resistance. **A** Heat map showing the loadings of all transcripts with loadings greater than 2.5 standard deviations from the mean in any tissue. The heat map was clustered using k medoid clustering. Functional enrichments of each cluster are indicated along the left margin. **B** Loadings for *Pparg* in different tissues. **C** Local and distal of *Pparg* expression in different tissues.

### Gene expression, but not local eQTLs, predicted body weight in an independent population

To test whether the transcript loadings identified in the DO could be translated to another population, we tested whether they could predict metabolic phenotype in an independent population of CC-RIX mice, which were F1 mice derived from multiple pairings of Collaborative Cross (CC) ^50;30;51;52^ strains (Fig. 7) (Methods). We tested two questions. First, we asked whether the loadings identified in the DO mice were relevant to the relationship between the transcriptome and the phenome in the CC-RIX. We predicted body weight (a surrogate for MDI) in each CC-RIX individual using measured gene expression in each tissue and the transcript loadings identified in the DO (Methods). The predicted body weight and acutal body weight were highly correlated (Fig. 7B left column). The best prediction was achieved for adipose tissue, which supports the observation in the DO that adipose expression was the strongest mediator of the genetic effect on MDI. This result also confirms the validity and translatability of the transcript loadings and their relationship to metabolic disease.

**Figure 7:**
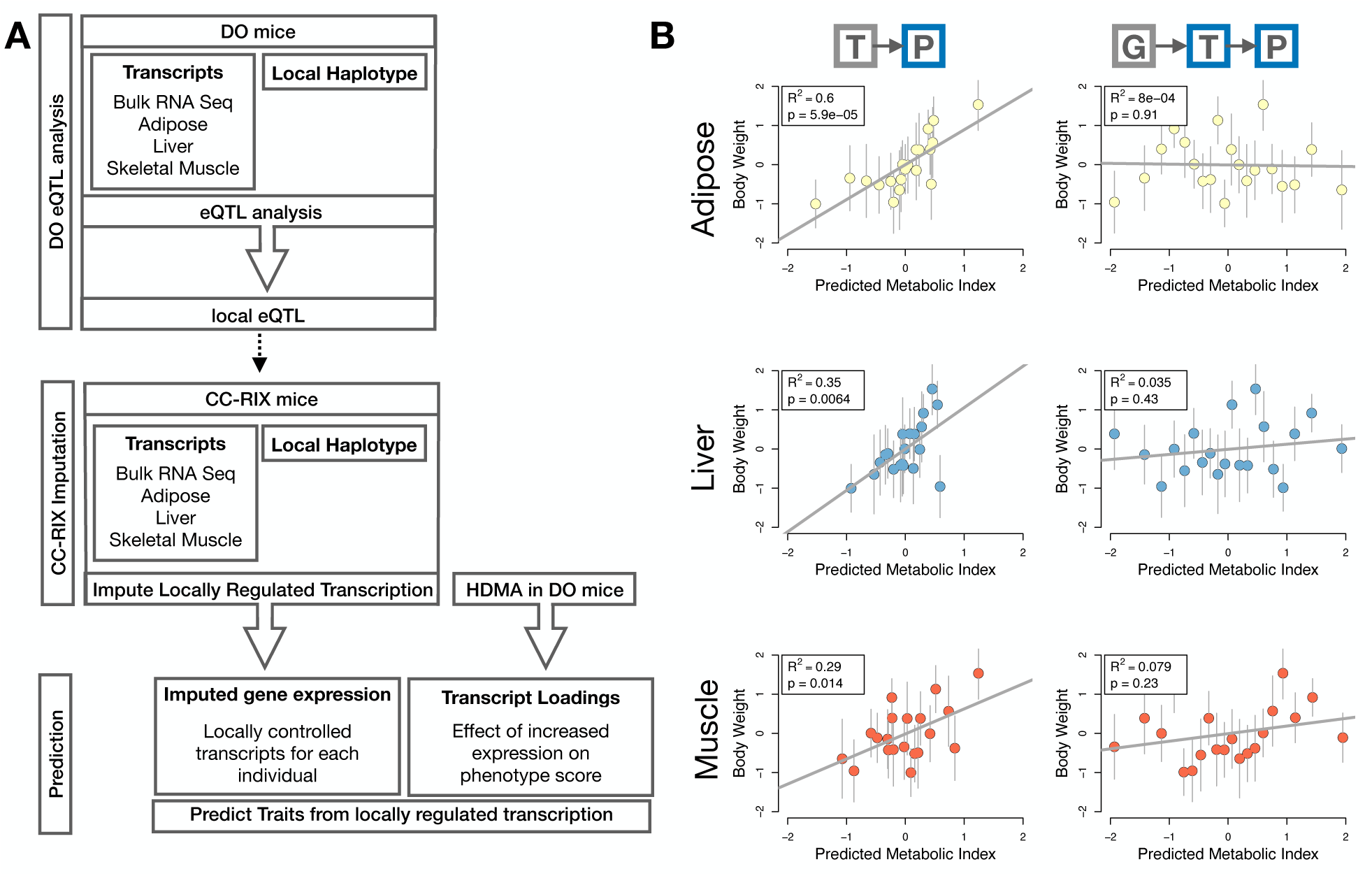
Transcription, but not local genotype, predicts phenotype in the CC-RIX. **A.** Workflow showing procedure for translating HDMA results to an independent population of mice. **B.** Relationships between the predicted metabolic disease index (MDI) and measured body weight. The left column shows the predictions using measured transcripts. The right column shows the prediction using transcript levels imputed from local genotype. Gray boxes indicate measured quantities, and blue boxes indicate calculated quantities. The dots in each panel represent individual CC-RIX strains. The gray lines show the standard deviation on body weight for the strain.

The second question related to the source of the relevant variation in gene expression. If local regulation was the predominant factor influencing trait-relevant gene expression, we should be able to predict phenotype in the CC-RIX using transcripts imputed from local genotype (Fig. 7A). The DO and the CC-RIX were derived from the same eight founder strains and so carry the same alleles throughout the genome. We imputed gene expression in the CC-RIX using local genotype and were able to estimate variation in gene transcription robustly (Supp. Fig. S8). However, these imputed values failed to predict body weight in the CC-RIX when weighted with the loadings from HDMA. (Fig. 7B right column). This result suggests that local regulation of gene expression is not the primary factor driving heritability of complex traits. It is also consistent with our findings in the DO population that distal heritability was a major driver of trait-relevant gene expression and that high-loading transcripts had comparatively high distal and low local heritability.

### Distally heritable transcriptomic signatures reflected variation in composition of adipose tissue and islets

The interpretation of global genetic influences on gene expression and phenotype is potentially more challenging than the interpretation and translation of local genetic influences, as genetic effects cannot be localized to individual gene variants or transcripts. However, there are global patterns across the loadings that can inform mechanism. For example, heritable variation in cell type composition can be inferred from transcript loadings. We observed above that immune activation in the adipose tissue was a highly enriched process correlating with obesity in the DO population. In humans, it has been extensively observed that macrophage infiltration in adipose tissue is a marker of obesity and metabolic disease ^53^. To determine whether the immune activation reflected a heritable change in cell composition in adipose tissue in DO mice, we compared loadings of cell-type specific genes in adipose tissue (Methods). The mean loading of macrophage-specific genes was significantly greater than 0 (Fig. 8A), indicating that obese mice were genetically predisposed to have high levels of macrophage infiltration in adipose tissue in response to the HFHS diet. Loadings for marker genes for other cell types were not statistically different from zero, indicating that changes in the abundance of those cell types is not a mediator of MDI.

**Figure 8:**
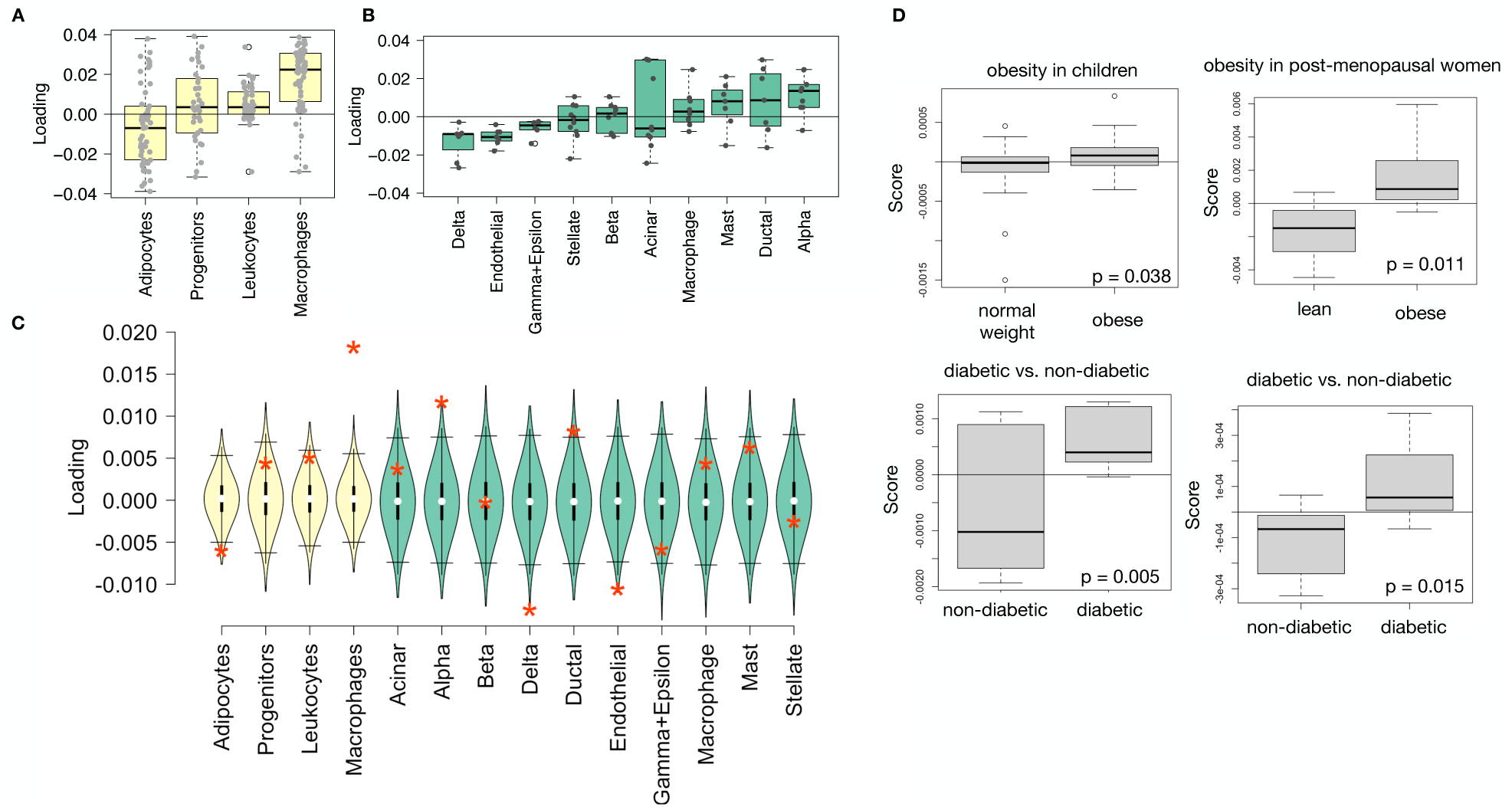
HDMA results translate to humans. **A.** Distribution of loadings for cell-type-specific transcripts in adipose tissue. **B.** Distribution of loadings for cell-type-specific transcripts in pancreatic islets (green). **C.** Null distributions for the mean loading of randomly selected transcripts in each cell type compared with the observed mean loading of each group of transcripts (red asterisk). **D.** Predictions of metabolic phenotypes in four adipose transcription data sets downloaded from GEO. In each study the obese/diabetic patients were predicted to have greater metabolic disease than the lean/non-diabetic patients based on the HDMA results from DO mice.

We also compared loadings of cell-type specific transcripts in islet (Methods). The mean loadings for alpha-cell specific transcripts were significantly greater than 0, while the mean loadings for delta- and endothelial-cell specific genes were significantly less than 0 (Fig. 8B). These results suggest that mice with higher MDI inherited an altered cell composition that predisposed them to metabolic disease, or that these compositional changes were induced by the HFHS diet in a heritable way. In either case, these results support the hypothesis that alterations in islet composition drive variation in MDI. Notably, the mean loading for pancreatic beta cell marker transcripts was not significantly different from zero. We stress that this is not necessarily reflective of the function of the beta cells in the obese mice, but rather suggests that any variation in the number of beta cells in these mice was unrelated to obesity and insulin resistance, the major contributors to MDI. This is further consistent with the islet composition traits having small loadings in the phenome score (Fig. 4).

### Heritable transcriptomic signatures translated to human disease

Ultimately, the heritable transcriptomic signatures that we identified in DO mice will be useful if they inform mechanism and treatment of human disease. To investigate the potential for translation of the gene signatures identified in DO mice, we compared them to transcriptional profiles in obese and non-obese human subjects (Methods). We limited our analysis to adipose tissue because the adipose tissue signature had the strongest relationship to obesity and insulin resistance in the DO.

We calculated a predicted MDI for each individual in the human studies based on their adipose tissue gene expression (Methods) and compared the predicted scores for obese and non-obese groups as well as diabetic and non-diabetic groups. In all cases, the predicted MDIs were higher on average for individuals in the obese and diabetic groups compared with the lean and non-diabetic groups (Fig. 8D). This indicates that the distally heritable signature of MDI identified in DO mice is relevant to obesity and diabetes in human subjects.

### Existing therapies are predicted to target mediator gene signatures

Another potential application of the transcript loading landscape is in ranking potential drug candidates for the treatment of metabolic disease. Although high-loading transcripts may be good candidates for understanding specific biology related to obesity, the transcriptome overall is highly interconnected and redundant. The ConnectivityMap (CMAP) database ^54^ developed by the Broad Institute allows querying thousands of compounds that reverse or enhance the extreme ends of transcriptomic signatures in multiple different cell types. By identifying drugs that reverse pathogenic transcriptomic signatures, we can potentially identify compounds that have favorable effects on gene expression. To test this hypothesis, we queried the CMAP database through the CLUE online query tool (https://clue.io/query/, version 1.1.1.43) (Methods). We identified top anti-correlated hits across all cell types (Supp. Figs S9 and S10). To get more tissue-specific results, we also looked at top results in cell types that most closely resembled our tissues. We looked at results in adipocytes (ASC) as well as pancreatic tumor cells (YAPC) regardless of *p* value (Supp. Figs S11 and S12).

Looking across all cell types, the notable top hits from the adipose tissue loadings included mTOR inhibitors and glucocorticoid agonists (Supp. Fig. S9). It is thought that metformin, which is commonly used to improve glycemic control, acts, at least in part, by inhibiting mTOR signaling ^55;56^. However, long-term use of other mTOR inhibitors, such as rapamycin, are known to cause insulin resistance and *β*-cell toxicity ^56–58^. Glucocorticoids are used to reduce inflammation, which was a prominent signature in the adipose tissue, but these drugs also promote hyperglycemia and diabetes ^59;60^. Accute treatment with glucocorticoids has further been shown to reduce thermogenesis in rodent adipocytes ^61–63^, but increase thermogenesis in human adipocytes ^64;65^. Thus, the pathways identified by CMAP across all cell types were highly related to the transcript loading profiles, but the relationship was not a simple reversal.

The top hit for the adipose composite transcript in CMAP adipocytes was a PARP inhibitor (Supp. Fig. S11). PARPs play a role in lipid metabolism and are involved in the development of obesity and diabetes ^66^. PARP1 inhibition increases mitochondrial biogenesis ^67^. Inihibition of PARP1 activity can further prevent necrosis in favor of the less inflammatory apoptosis ^68^, thereby potentially reducing inflammation in stressed adipocytes. Other notable hits among the top 20 were BTK inhibitors, which have been observed to suppress inflammation and improve insulin resistance ^69^ as well as to reduce insulin antibodies in type I diabetes ^70^. IkappaB kinase (IKK) is an enzyme complex involved in regulating cellular responses to inflammation ^71^. Inhibitors of IKK have been shown to improve glucose control in type II diabetes ^72;73^.

Among the top most significant hits for the transcript loadings from pancreatic islets (Supp. Fig. S10), was suppression of T cell receptor signaling, which is known to be involved in Type 1 diabetes ^74^, as well as TNFR1, which has been associated with mortality in diabetes patients ^75^. Suppression of NOD1/2 signaling was also among the top hits. NOD1 and 2 sense ER stress ^76;77^, which is associated with *β*-cell death in type 1 and type 2 diabetes ^78^. This cell death process is dependent on NOD1/2 signaling ^76^, although the specifics have not yet been worked out.

We also looked specifically at hits in pancreatic tumor cells (YAPC) regardless of significance level to get a transcriptional response more specific to the pancreas (Supp. Fig. S12). Hits in this list included widely used diabetes drugs, such as sulfonylureas, PPAR receptor agonists, and insulin sensitizers. Rosiglitazone is a PPAR-*γ* agonist and was one of the most prescribed drugs for type 2 diabetes before its use was reduced due to cardiac side-effects ^79^. Sulfonylureas are another commonly prescribed drug class for type 2 diabetes, but also have notable side effects including hypoglycemia and accellerated *β*-cell death ^80^.

In summary, the high-loading transcripts derived from HDMA in mice prioritized of drugs with demonstrated effectiveness in reducing type 2 diabetes phenotypes in humans in a tissue-specific manner. Drugs identified using the islet loadings are known diabetes drugs that act directly on pancreatic function. Drugs identified by the adipose loadings tended to reduce inflammatory responses and have been shown incidentally to reduce obesity-related morbidity.

## Discussion

Here we investigated the relative contributions of local and distal gene regulation in four tissues to heritable variation in traits related to metabolic disease in genetically diverse mice. We found that distal heritability was positively correlated with trait relatedness, whereas high heritability was negatively correlated with trait relatedness. We used a novel high-dimensional mediation analysis (HDMA) to identify tissue-specific composite transcripts that are predicted to mediate the effect of genetic background on metabolic traits. The adipose-derived composite transcript robustly predicted body weight in an independent cohort of diverse mice with disparate population structure, as well as to humans. However, gene expression imputed from local genotype failed to predict body weight in the second population. Taken together, these results highlight the complexity of gene expression regulation in relation to trait heritability and suggest that heritable trait variation is mediated primarily through distal gene regulation.

## Supporting information

Tissue of action gene list

## Supplemental Discussion

Our result that distal regulation accounted for most trait-related gene expression differences is consistent with a complex model of genetic trait determination. It has frequently been assumed that gene regulation in *cis* is the primary driver of genetically associated trait variation, but attempts to use local gene regulation to explain phenotypic variation have had limited success ^16;17^. In recent years, evidence has mounted that distal gene regulation may be an important mediator of trait heritability ^19;18;81^. It has been observed that transcripts with high local heritability explain less expression-mediated disease heritability than those with low local heritability ^19^. Consistent with this observation, genes located near GWAS hits tend to be complexly regulated ^18^. They also tend to be enriched with functional annotations, in contrast to genes with simple local regulation, which tend to be depleted of functional annotations suggesting they are less likely to be directly involved in disease traits ^18^. These observations are consistent with principles of robustness in complex systems in which simple regulation of important elements leads to fragility of the system ^82–84^. Our results are consistent, instead, with a more complex picture where genes whose expression can drive trait variation are buffered from local genetic variation but are extensively influenced indirectly by genetic variation in the regulatory networks converging on those genes.

Our results are also consistent with the recently proposed omnigenic model, which posits that complex traits are massively polygenic and that their heritability is spread out across the genome ^85^. In the omnigenic model, genes are classified either as “core genes,” which directly impinge on the trait, or “peripheral genes,” which are not directly trait-related, but influence core genes through the complex gene regulatory network. HDMA explicitly models a central proposal of the omnigenic model which posits that once the expression of the core genes (i.e. trait-mediating genes) is accounted for, there should be no residual correlation between the genome and the phenome. Here, we were able to fit this model and identified a composite transcript that, when taken into account, left no residual correlation between the composite genome and composite phenome scores (Fig. 3A).

Unlike in the omnigenic model, we did not observe a clear demarcation between the core and peripheral genes in loading magnitude, but we do not necessarily expect a clear separation given the complexity of gene regulation and the genotype-phenotype map ^86^. An extension of the omnigenic model proposed that most heritability of complex traits is driven by weak distal eQTLs that are potentially below the detection threshold in studies with feasible sample sizes ^81^. This is consistent with what we observed here. For example, *Nucb2*, had a high loading in islets and was also strongly distally regulated (66% distal heritability) (Fig. 5). This gene is expressed in pancreatic *β* cells and is involved in inslin and glucagon release ^87–89^. Although its transcription was highly heritable in islets, that regulation was distributed across the genome, with no clear distal eQTL (Supp. Fig. S13). Thus, although distal regulation of some genes may be strong, this regulation is likely to be highly complex and not easily localized.

Individual high-loading transcripts also demonstrated biologically interpretable, tissue-specific patterns. We highlighted *Pparg*, which is known to be protective in adipose tissue ^44^ where it was negatively loaded, and harmful in the liver ^45–49^, where it was positively loaded. Such granular patterns may be useful in generating hypotheses for further testing, and prioritizing genes as therapeutic targets. The tissue-specific nature of the loadings also may provide clues to tissue-specific effects, or side effects, of targeting particular genes system-wide.

In addition to identifying individual transcripts of interest, the composite transcripts can be used as weighted vectors in multiple types of analysis, such as drug prioritization using gene set enrichment analysis (GSEA) and the CMAP database. In particular, the CMAP analysis identified drugs which have been demonstrated to reverse insulin resistance and other aspects of metabolic disease. This finding supports the causal role of these full gene signatures in pathogenesis of metabolic disease and thus their utility in prioritizing drugs and gene targets as therapeutics.

Together, our results have shown that both tissue specificity and distal gene regulation are critically important to understanding the genetic architecture of complex traits. We identified important genes and gene signatures that were heritable, plausibly causal of disease, and translatable to other mouse populations and to humans. Finally, we have shown that by directly acknowledging the complexity of both gene regulation and the genotype-to-phenotype map, we can gain a new perspective on disease pathogenesis and develop actionable hypotheses about pathogenic mechanisms and potential treatments.

## Data and Code Availability

**DO mice:** Genotypes, phenotypes, and pancreatic islet gene expression data were previously published ^12^. Gene expression for the other tissues can be found at the Gene Expression Omnibus https://www.ncbi.n lm.nih.gov/geo/ with the following accession numbers: DO adipose tissue - GSE266549; DO liver tissue - GSE266569; DO skeletal muscle - GSE266567. Expression data with calculated eQTLs are available at Figshare https://figshare.com/ DOI: 10.6084/m9.figshare.27066979

**CC-RIX mice**: Gene expression can be found at the Gene Expression Omnibus https://www.ncbi.nlm.nih.gov/geo/ with the following accession numbers: CC-RIX adipose tissue - GSE237737; CC-RIX liver tissue - GSE237743; CC-RIX skeletal muscle - GSE237747. Count matrices and phenotype data can be found at Figshare https://figshare.com/ DOI: 10.6084/m9.figshare.27066979

**Code**: All code used to run the analyses reported here are available at Figshare: https://figshare.com/ DOI: 10.6084/m9.figshare.27066979

## Acknowledgements

This project was supported by The Jackson Laboratory Cube Initiative, as well as grants from the National Institutes of Health (grant numbers.: R01DK101573, R01DK102948, and RC2DK125961) (to A. D. A.) and by the University of Wisconsin-Madison, Department of Biochemistry and Office of the Vice Chancellor for Research and Graduate Education with funding from the Wisconsin Alumni Research Foundation (to M. P. K.).

We thank the following scientific services at The Jackson Laboratory: Genome Technologies for the RNA sequencing, necropsy services for the tissue harvests, and the Center for Biometric Analysis for metabolic phenotyping.

## Supplemental Figures

**Figure S1:**
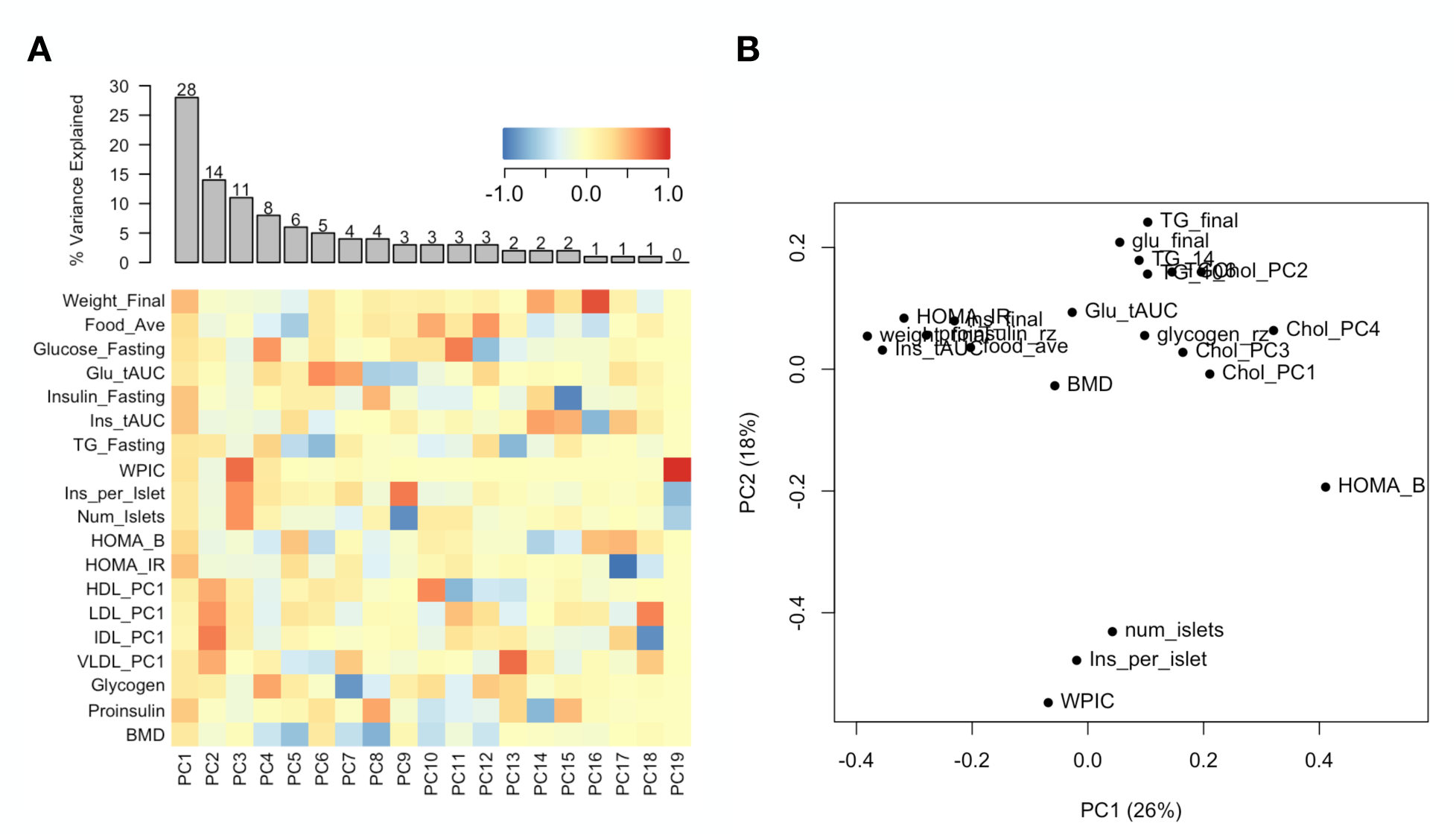
Trait matrix decomposition. **A** The heat map shows the loadings of each trait onto each principal component of the trait matrix. The bars at the top show the percent variance explained for each principal component. **B** Traits plotted by the first and second principal components of the trait matrix. This view shows clustering of traits into insulin- and weight-related traits, lipid-related traits, and ex-vivo pancreatic measurements.

**Figure S2:**
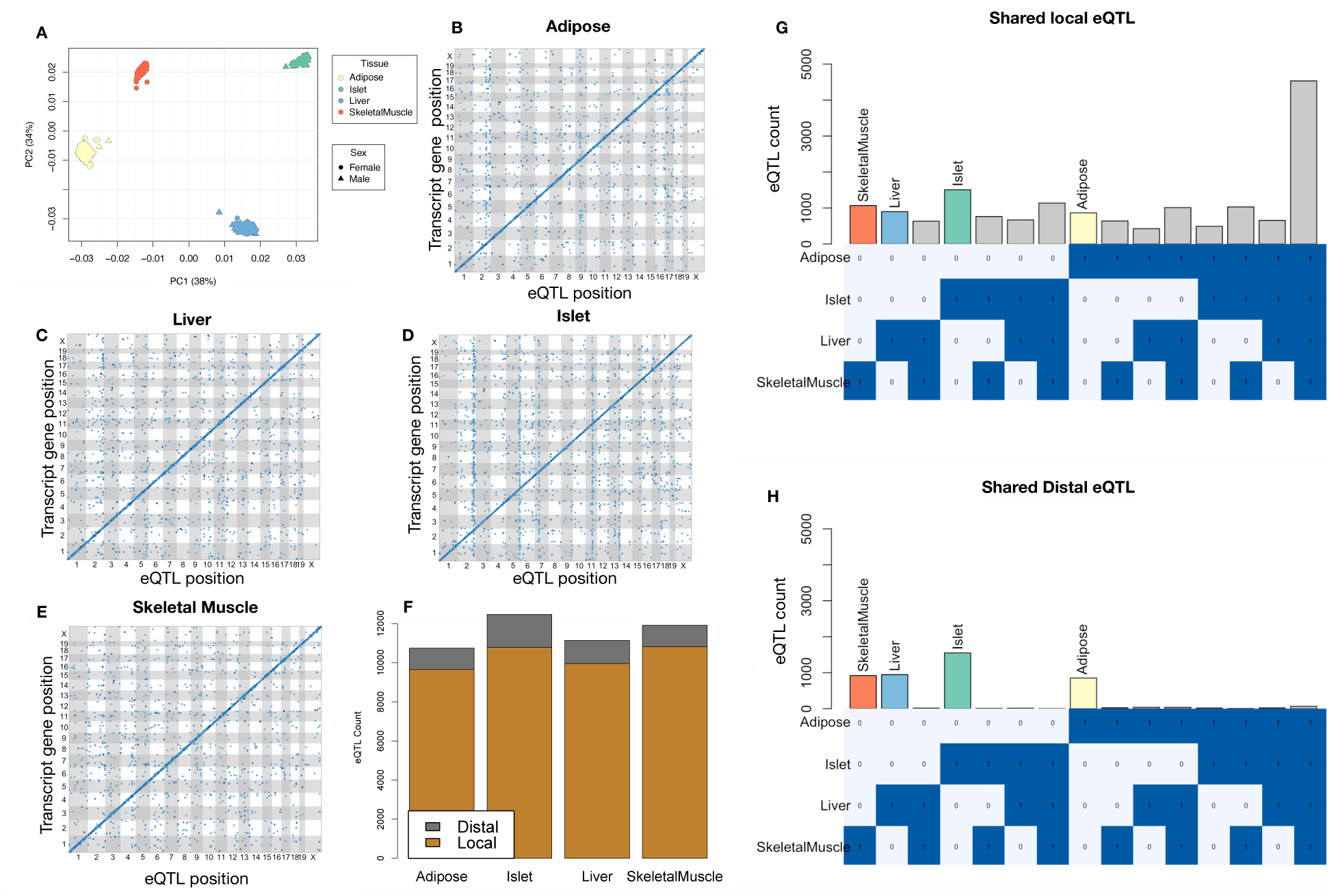
Overview of eQTL analysis in DO mice. **A.** RNA seq samples from the four different tissues clustered by tissue. **B.-E.** eQTL maps are shown for each tissue. The *x*-axis shows the position of the mapped eQTL, and the *y*-axis shows the physical position of the gene encoding each mapped transcript. Each dot represents an eQTL with a minimum LOD score of 8. The dots on the diagonal are locally regulated eQTL for which the mapped eQTL is at the within 4Mb of the encoding gene. Dots off the diagonal are distally regulated eQTL for which the mapped eQTL is distant from the gene encoding the transcript. **F.** Comparison of the total number of local and distal eQTL with a minimum LOD score of 8 in each tissue. All tissues have comparable numbers of eQTL. Local eQTLs are much more numerous than distal eQTL. **G.** Counts of transcripts with local eQTL shared across multiple tissues. The majority of local eQTLs were shared across all four tissues. **H.** Counts of transcripts with distal eQTL shared across multiple tissues. The majority of distal eQTL were tissue-specific and not shared across multiple tissues. For both G and H, eQTL for a given transcript were considered shared in two tissues if they were within 4Mb of each other. Colored bars indicate the counts for individual tissues for easy of visualization.

**Figure S3:**
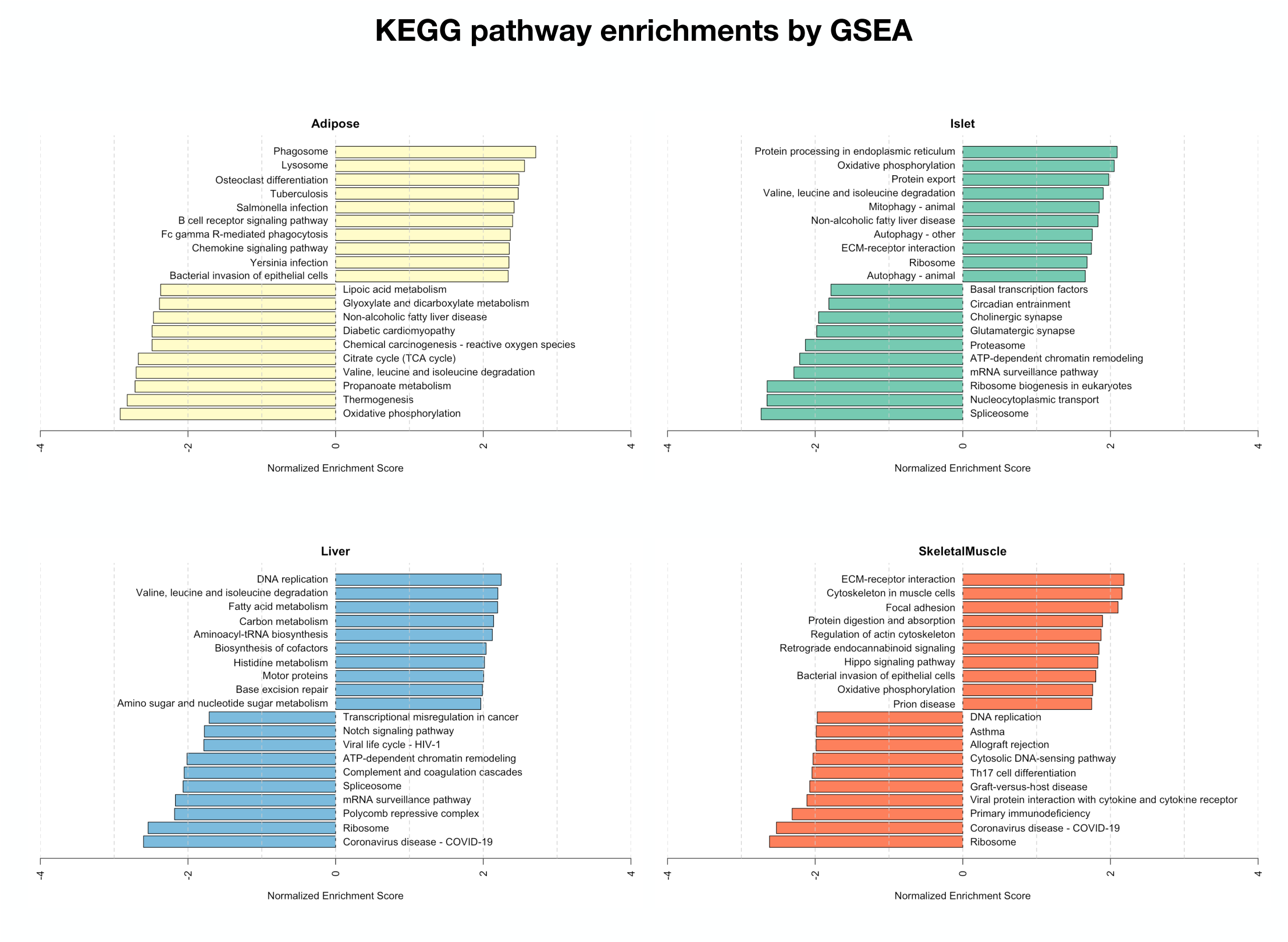
Bar plots showing normalized enrichment scores (NES) for KEGG pathways as determined by fast gene score enrichment analysis (fgsea). Only the top 10 positive and top 10 negative scores are shown. Colors indicate tissue. The name beside each bar shows the name of each enriched KEGG pathway.

**Figure S4:**
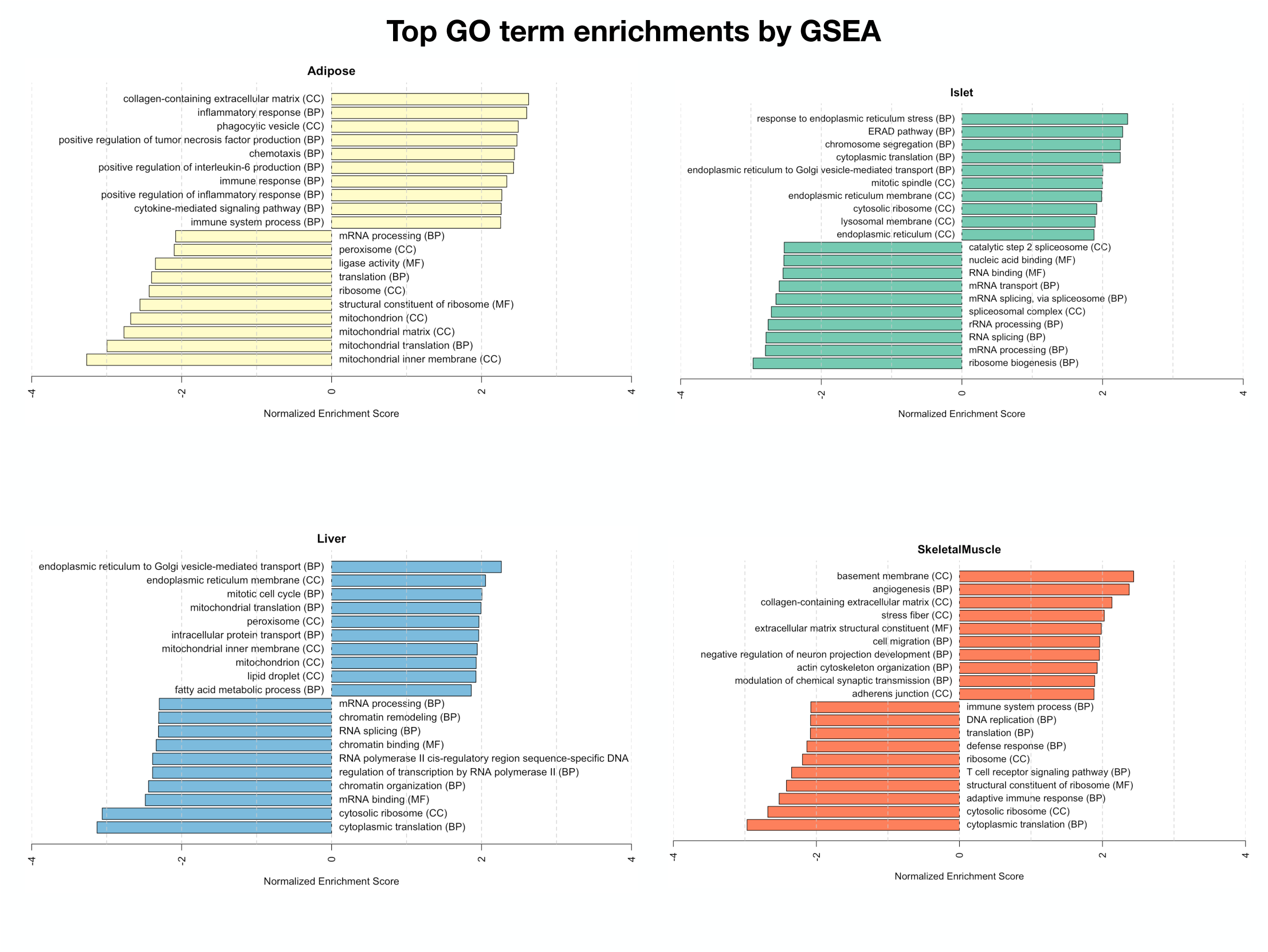
Bar plots showing normalized enrichment scores (NES) for GO terms as determined by fast gene score enrichment analysis (fgsea). Only the top 10 positive and top 10 negative scores are shown. Colors indicate tissue. The name beside each bar shows the name of each enriched GO term. The letters in parentheses indicate whether the term is from the biological process ontology (BP), the molecular function ontology (MF), or the cellular compartment ontology (CC).

**Figure S5:**
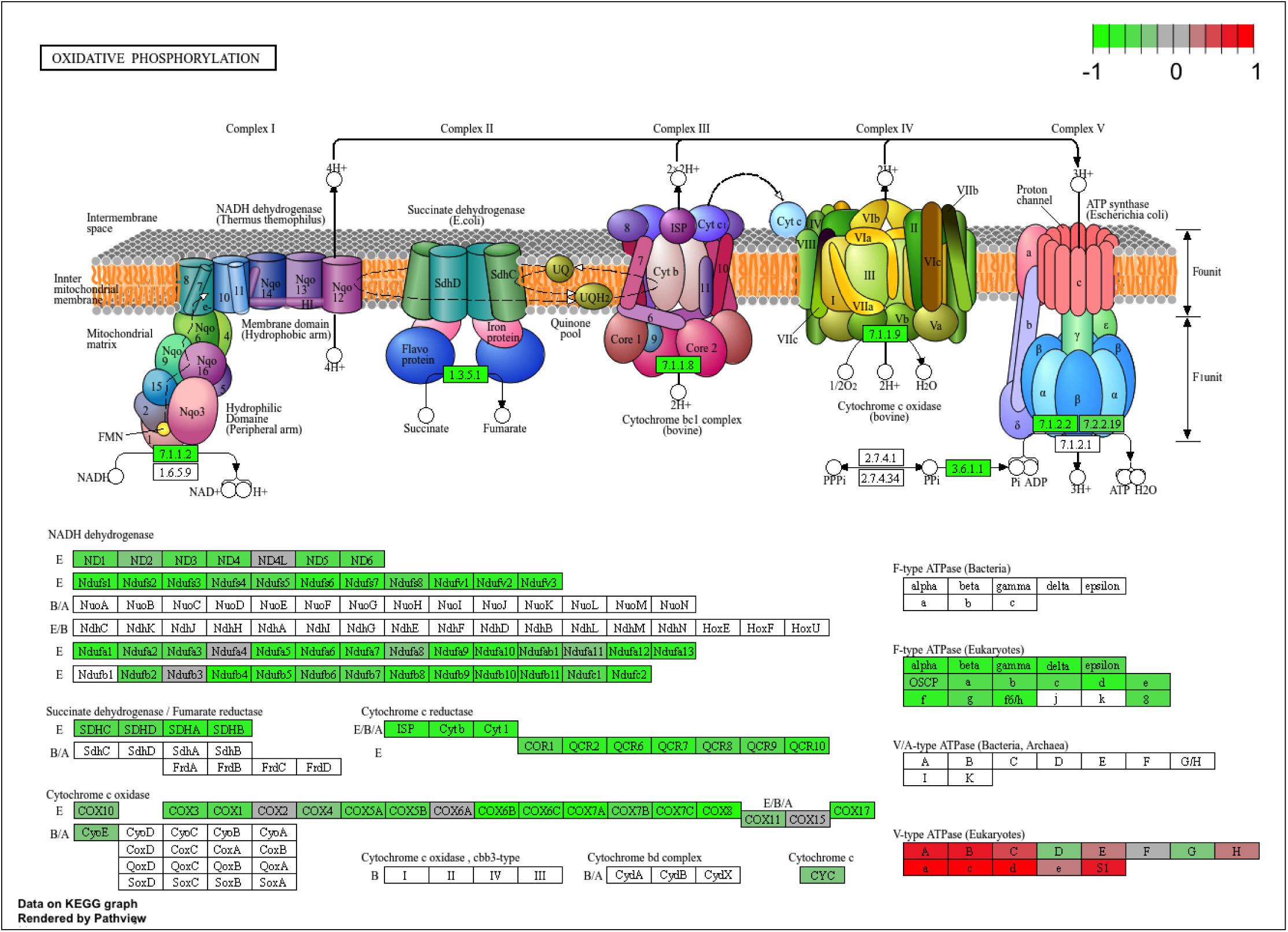
The KEGG pathway for oxidative phosphorylation in mice. Each element is colored based on its HDMA loading from adipose tissue normalized to run from -1 to 1. Genes highlighted in green had negative loadings, and those highlighted in red had positive loadings. Almost the entire pathway was strongly negatively loaded indicating that increased expression of genes involved in oxidative phosphorylation was associated with reduced MDI.

**Figure S6:**
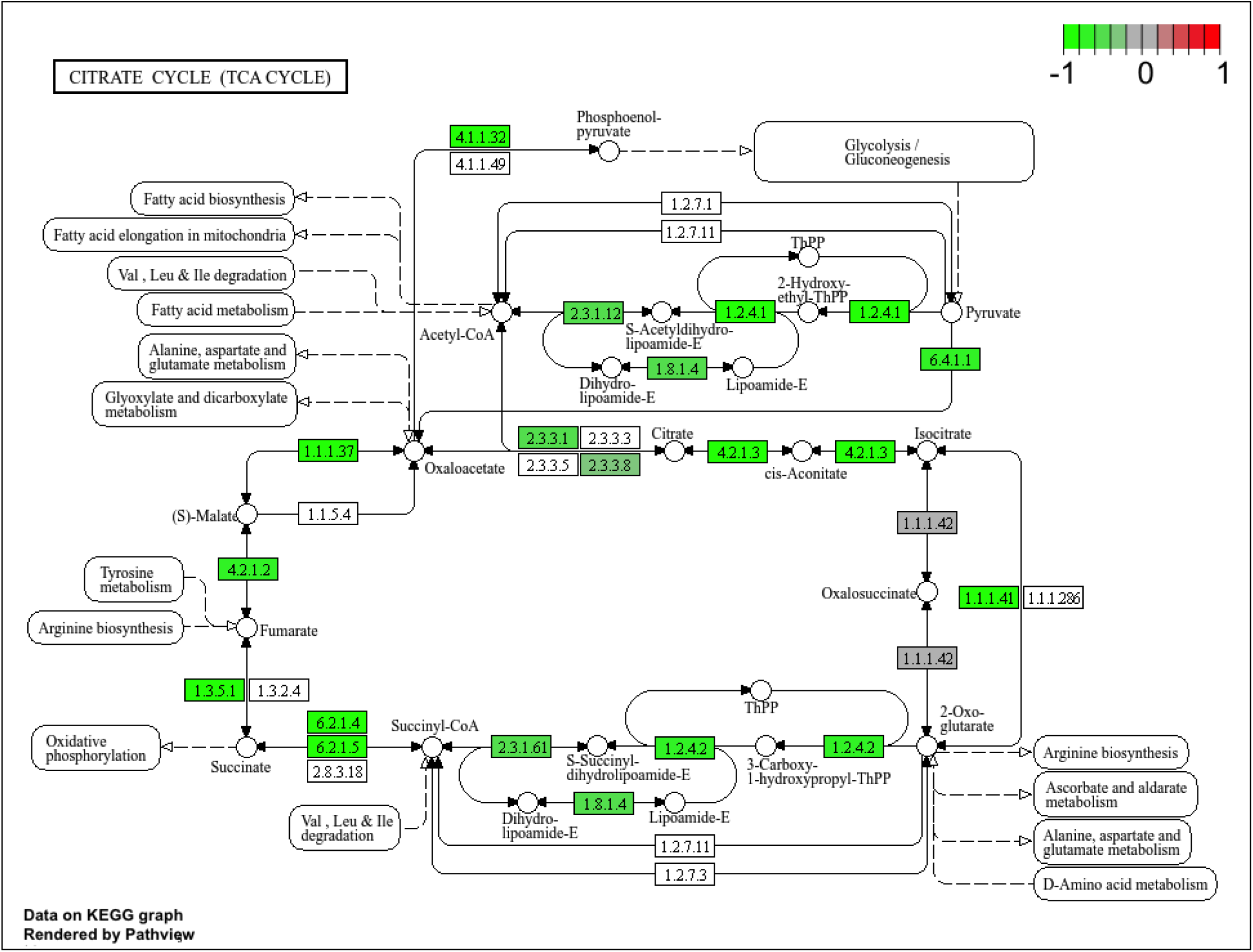
The KEGG pathway for the TCA (citric acid) cycle in mice. Each element is colored based on its HDMA loading from adipose tissue normalized to run from -1 to 1. Genes highlighted in green had negative loadings, and those highlighted in red had positive loadings. Many genes in the cycle were strongly negatively loaded indicating that increased expression of genes involved in branched-chain amino acid degradation was associated with reduced MDI.

**Figure S7:**
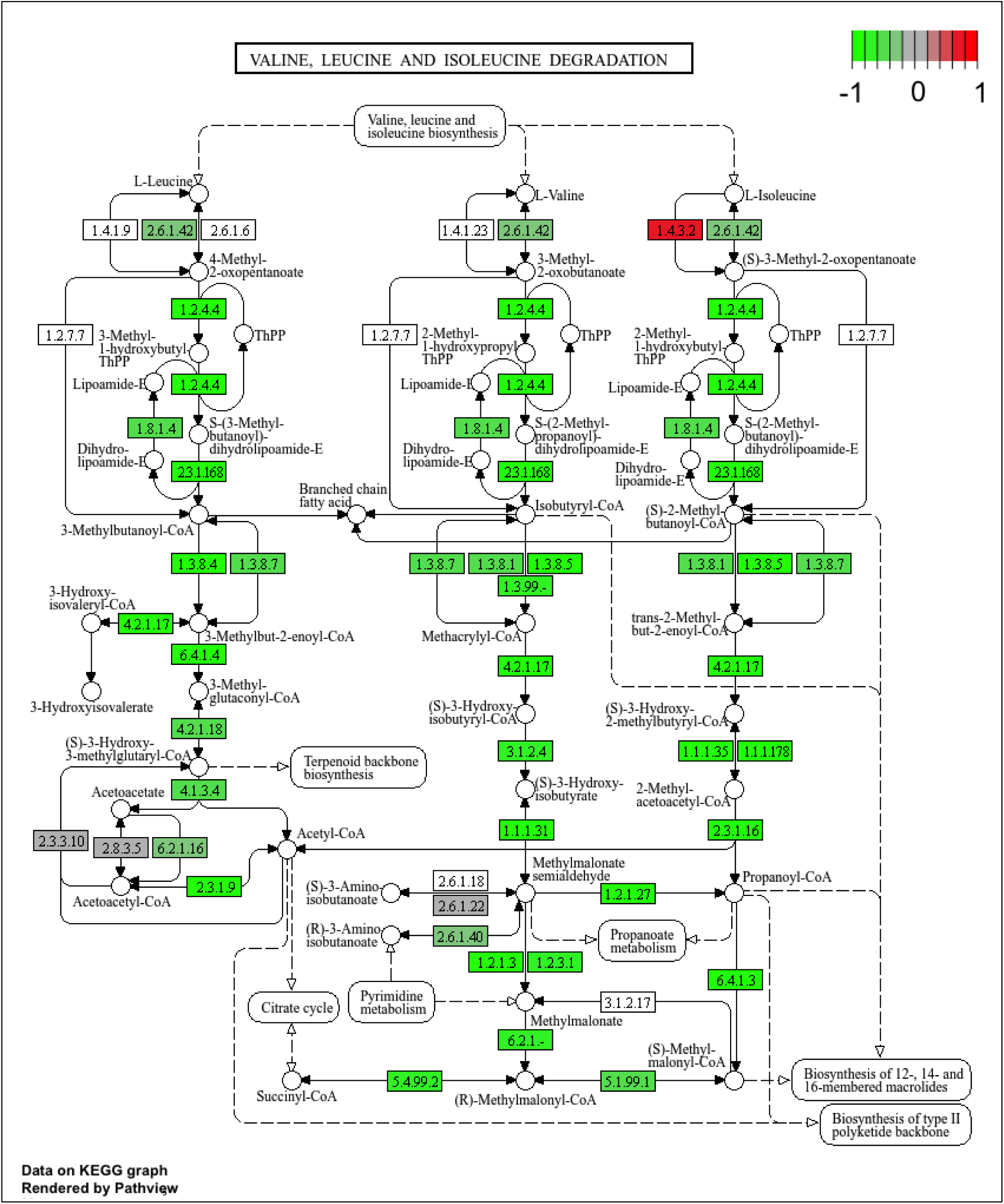
The KEGG pathway for branched-chain amino acid degradation in mice. Each element is colored based on its HDMA loading from adipose tissue normalized to run from -1 to 1. Genes highlighted in green had negative loadings, and those highlighted in red had positive loadings. Almost the entire pathway was strongly negatively loaded indicating that increased expression of genes involved in branched-chain amino acid degradation was associated with reduced MDI.

**Figure S8:**
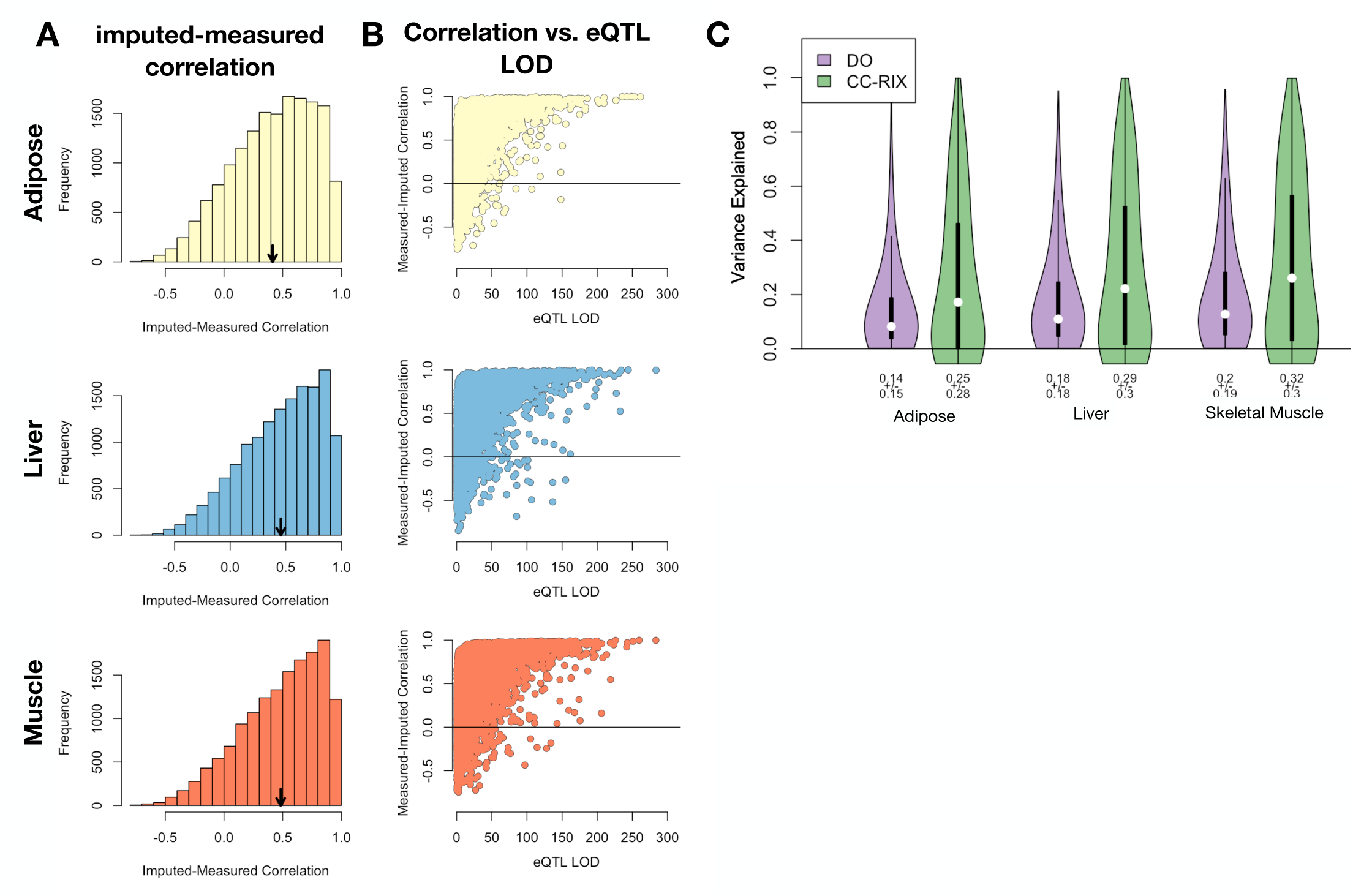
Validation of transcript imputation in the CC-RIX. **A.** Distributions of correlations between imputed and measured transcripts in the CC-RIX. The mean of each distribution is shown by the red line. All distributions were skewed toward positive correlations and had positive means near a Pearson correlation (r) of 0.5. **B.** The relationship between the correlation between measured and imputed expression in the CC-RIX (x-axis) and eQTL LOD score. As expected, imputations are more accurate for transcripts with strong local eQTLs. **C.** Variance explained by local genotype in the DO and CC-RIX.

**Figure S9:**
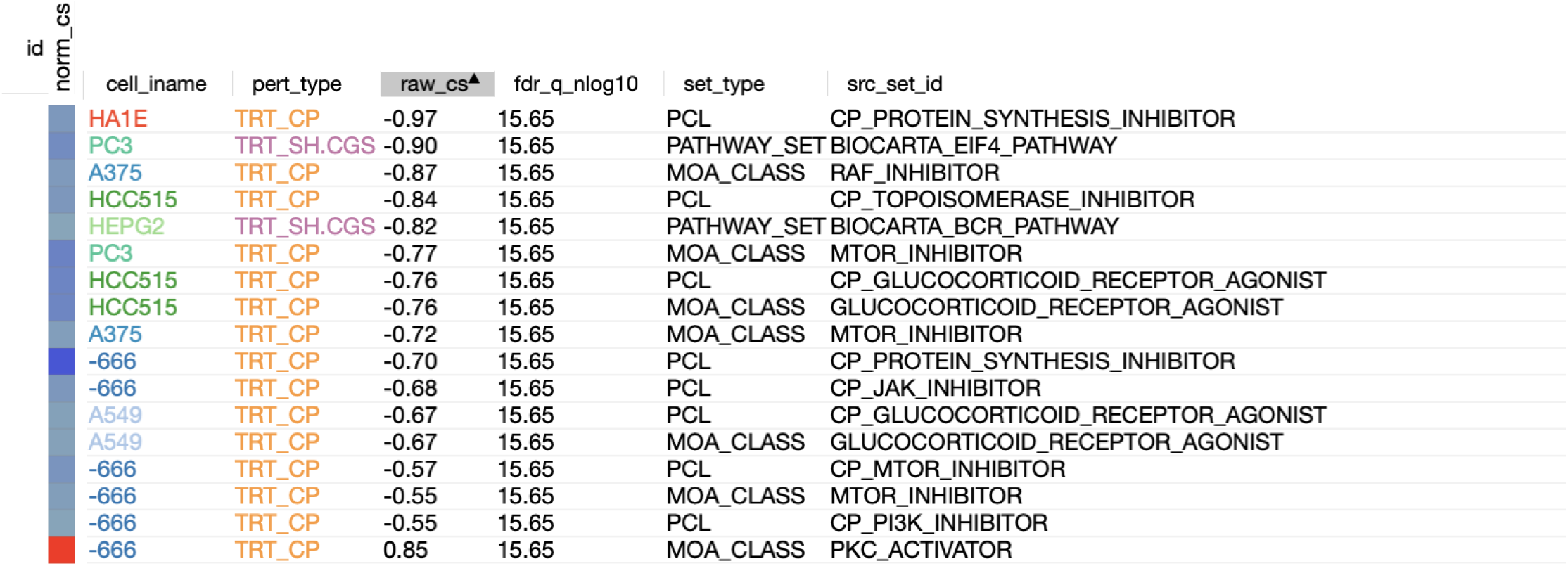
CMAP results using the *adipose* tissue composite transcript as an input. Table includes results from *all cell types* sorted with a *−log*_10_(*q*) *>* 15. The results are sorted by the correlation of the query to the input with the most negative results at the top.

**Figure S10:**
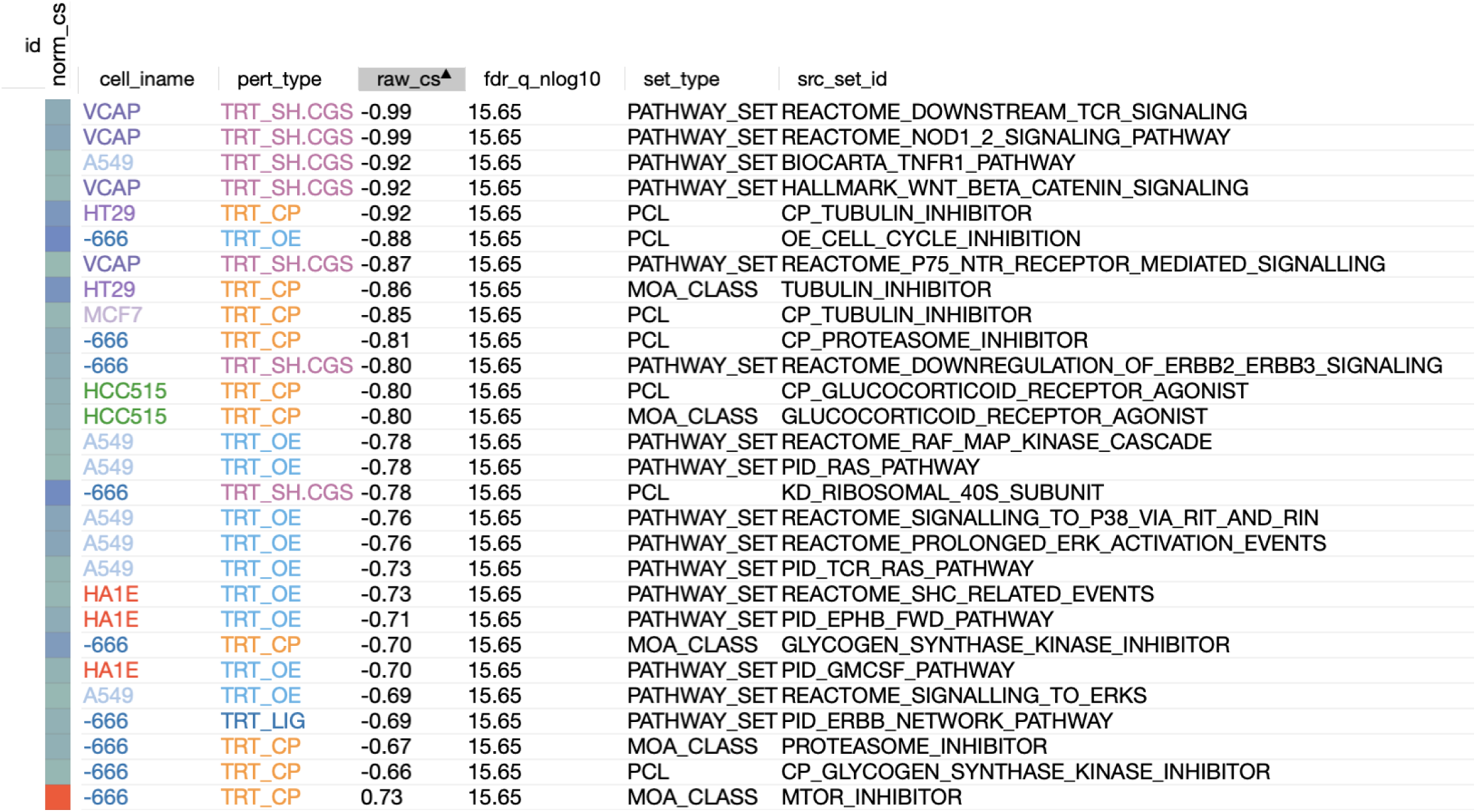
CMAP results using the *pancreatic islet* tissue composite transcript as an input. Table includes results from *all cell types* sorted with a *−log*_10_(*q*) *>* 15. The results are sorted by the correlation of the query to the input with the most negative results at the top.

**Figure S11:**
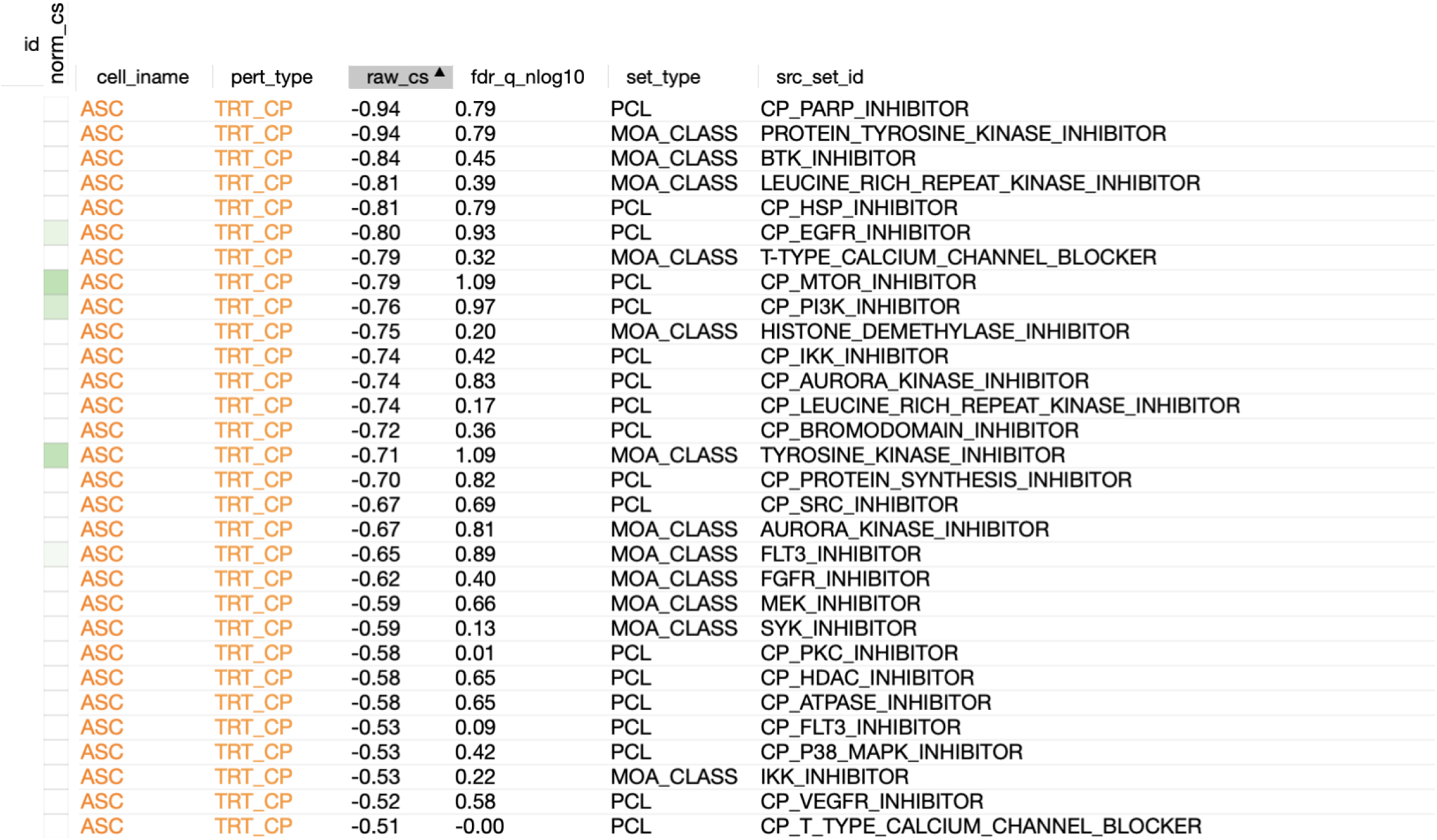
CMAP results using the *adipose* tissue composite transcript as an input. Table includes the top 30 results derived *only from normal adipocytes* (ASC) regardless of significance. The results are sorted by the correlation of the query to the input with the most negative results at the top.

**Figure S12:**
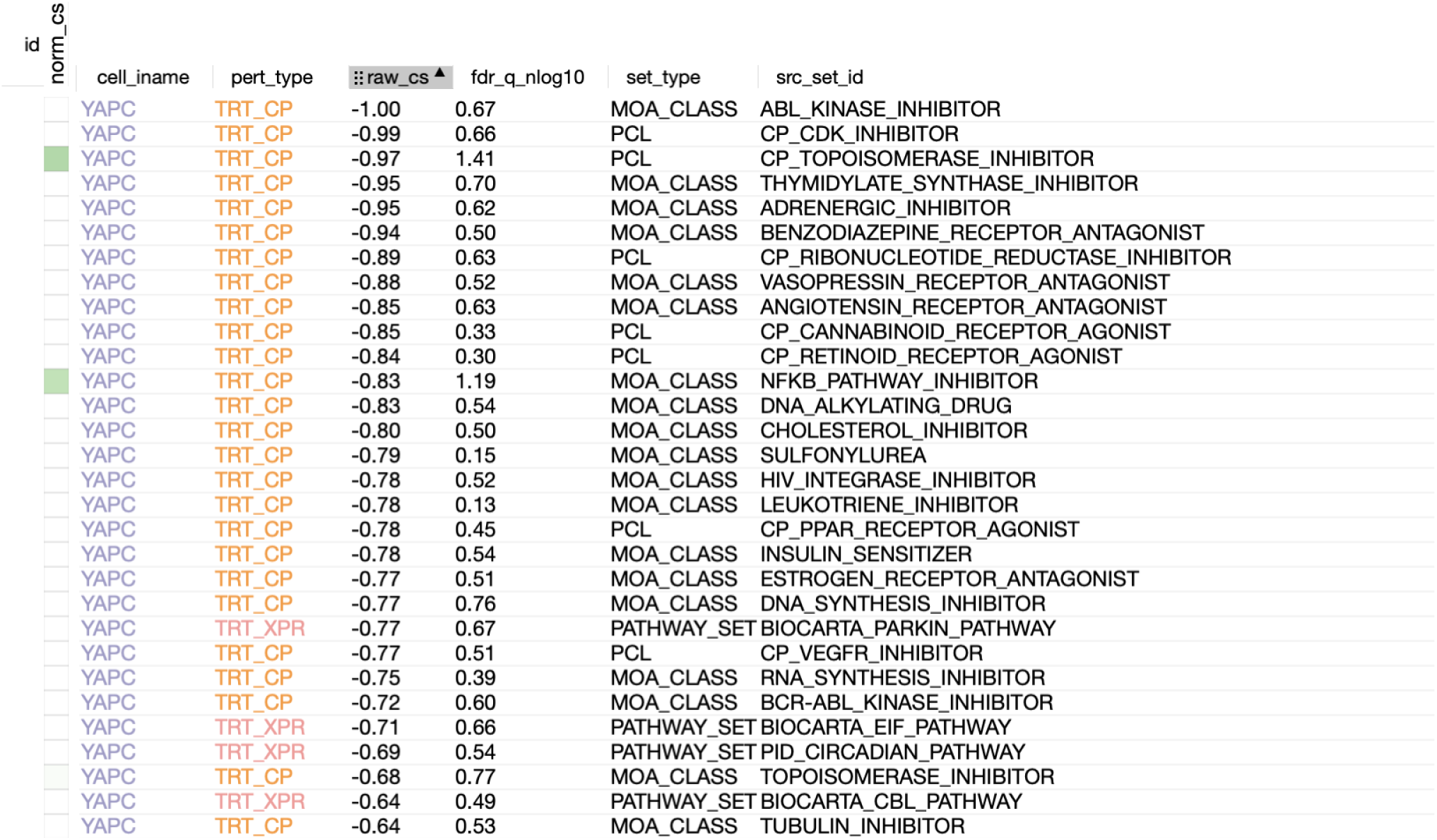
CMAP results using the *pancreatic islet* composite transcript as an input. Table includes the top 30 results derived *only from YAPC cells*, which are derived from pancreatic carcinoma cells. Results are shown regardless of significance and are sorted by the correlation of the query to the input with the most negative results at the top.

**Figure S13:**
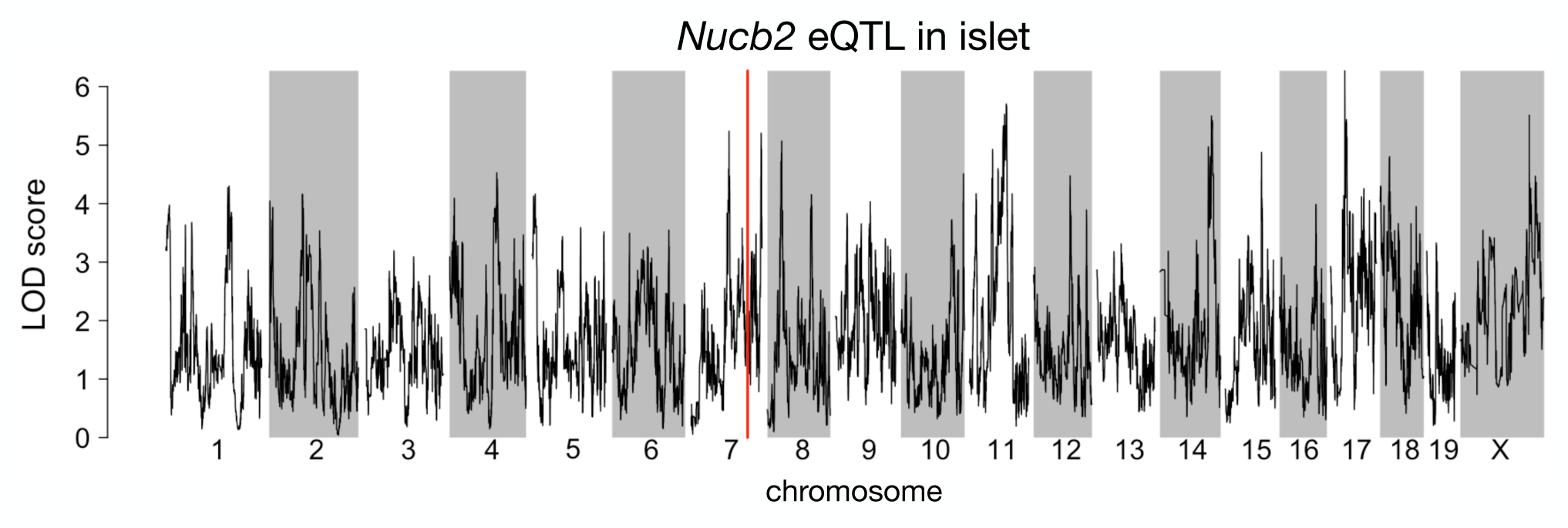
Regulation of *Nucb2* expression in islet. *Nucb2* is encoded on mouse chromosome 7 at 116.5 Mb (red line). In islets the heritability of *Nucb2* expression levels is 69% heritable. This LOD score trace shows that there is no local eQTLs at the position of the gene, nor any strong distal eQTL anywhere else in the genome.

## Notes

### Competing Interest Statement

The authors have declared no competing interest.

