## Supplementary material for "Transcripts with high distal heritability mediate genetic effects on complex metabolic traits": Online Methods

### Online methods for: Transcripts with high distal heritability mediate genetic effects on complex metabolic traits

#### Diversity Outbred Mice

Mice were maintained and treated in accordance with the guidelines approved by the Department of Biochemistry animal vivarium at the University of Wisconsin. Animal husbandry and in vivo phenotyping methods were previously published[1;2].

A population of 500 diversity outbred mice (split evenly between male and female) from generates 18, 19, and 21, was placed on a high-fat (44.6% kcal fat), high-sugar (34% carbohydrate), adequate protein (17.3 % protein) diet from Envigo Teklad (catalog number TD.08811) starting at four weeks of age as described previously[2]. Individuals were assessed longitudinally for multiple metabolic measures including fasting glucose levels, glucose tolerance, insulin levels, body weight, and blood lipid levels.

#### Trait measurements

Trait measurements were described previously in[2]. Briefly, body weight was measured every two weeks, and 4-hour fasting plasma samples were collected to measure insulin, glucose, and triglycerides (TG). At around 18 weeks of age, an oral glucose tolerance test (oGTT) was conducted on 4-hour fasted mice to assess changes in plasma insulin and glucose. Glucose (2 g/kg) was given via oral gavage. Blood samples were taken from a retro-orbital bleed before glucose administration, and at 5, 15, 30, 60, and 120 minutes afterward. The area under the curve (AUC) was calculated for glucose and insulin. Glucose was measured using the glucose oxidase method, and insulin was measured by radioimmunoassay.

HOMA-IR and HOMA-B, which are homeostatic model assessments of insulin resistance (IR) and pancreatic islet function (B), were calculated using fasting plasma glucose and insulin values at the start of the oGTT. HOMA-IR = (glucose $\times$ insulin) / 405 and HOMA-B = (360 $\times$ insulin) / (glucose - 63). Plasma glucose and insulin units are mg/dL and mU/L, respectively.

#### Genotyping

Genotypes at 143,259 markers was performed using the Mouse Universal Genotyping Array (GigaMUGA)[3] at Neogen (Lincoln, NE) as described previously[2;4]. Genotypes were converted to founder strain-haplotype reconstructions using the R/DOQTL software[5] and interpolated onto a grid with 0.02-cM spacing to yield 69,005 pseudomarkers. Individual chromosome (Chr) haplotypes were reconstructed from RNA-seq data using a hidden Markov model (GBRS, https://github.com/churchill-lab/gbrs).

#### Processed Data

We downloaded genotypes, phenotypes, and pancreatic islet gene expression data from Dryad (doi:10.5061/dryad.pj105).

#### Collaborative cross recombinant inbred mice

Mice were cared for and treated following the guidelines approved by the Association for Assessment and Accreditation of Laboratory Animal Care at The Jackson Laboratory. All animals were obtained from The Jackson Laboratory. The mice were kept in a pathogen-free room at a temperature ranging from 20 to 22°C with a 12-hour light/dark cycle. Starting at 6 weeks of age, they were fed either a custom-designed high-fat, high-sugar (HFHS) diet (Research Diets D19070208) or a control diet (Research Diets D19072203) *ad libitum*. Body weight was measured weekly until the mice were about 16 weeks old, after which measurements were taken every other week. Food intake measurements were collected at 14 weeks, 23 weeks (for 6-month cohorts), 26 weeks (for 12-month cohorts), 38 weeks, and 51 weeks by weighing the grain contents in the cage over a three-day period. Fasted serum was collected at 14 weeks, 28 weeks (for 6-month cohorts), 26 weeks (for 12-month cohorts), 38 weeks, and 56 weeks of age via retro-orbital or submental vein. In a subset of the 12-month cohort, metformin (5 mg/mL) was administered *ad libitum* in the drinking water. The first two weeks of treatment included monitoring water intake to ensure consumption. Metformin treatment continued for the duration of the experiment.

#### Clinical chemistries

CC-RIX animals were fasted for four hours before serum collection via the retro-orbital or submental vein. Whole blood was left at room temperature for 30-60 minutes before being centrifuged for 5 minutes at 12,500 RPM. The serum was then tested for glucose (Beckman Coulter; OSR6121), cholesterol (Beckman Coulter; OSR6116), triglycerides (Beckman Coulter; OSR60118), insulin (MSD; K152BZC-1), or c-peptide (MSD; K1526JK-1).

#### Intraperitoneal glucose tolerance testing

After a fasting period of 4-6 hours, baseline glucose measurements were taken from CC-RIX mice using an AlphaTrak2 glucometer and test strips (Zoetis) by making a small nick in the tail tip. A bolus intraperitoneal injection of 20% glucose (1g/kg) was then administered, and additional tail tip nicks were performed at 15, 30, 60, and 120 minutes post-injection to measure glucose levels.

#### Dual Energy X-ray Absorptiometry (DEXA)

To assess bone mineral density in the CC-RIX population at either 27 weeks of age (6-month cohorts) or 55 weeks of age (12-month cohorts), the mice were weighed and anesthetized through continuous inhalation of isoflurane. The Faxitron UltraFocus DXA system was used to emit two energy levels, 40 kV and 80 kV, for capturing images of bone and soft tissue.

#### Bulk tissue collection

At either 28 weeks of age (for the 6-month cohort) or 56 weeks of age (for the 12-month cohort), CC-RIX animals were humanely euthanized by cervical dislocation. Tissues, including adipose, gastrocnemius, and the left liver lobe, were harvested and flash-frozen in liquid nitrogen for RNA sequencing.

#### Whole Pancreas Insulin Content

The animals were humanely euthanized at 16 weeks of age and the entire pancreas was removed, ensuring no excess fat or mesentery tissue was included. The pancreas tissue was placed in a pre-weighed 20 mL glass scintillation vial containing acid ethanol (75% HPLC grade ethanol (ThermoFisher; A995-4), 1.\5% concentrated hydrochloric acid (ThermoFisher; A144-212) in distilled water). The weight of the pancreas was measured for normalization. Using curved scissors, the pancreas was chopped for four minutes, and the samples were stored at $-20°$C until all animals were harvested. For insulin measurements, the contents of the scintillation vials were rinsed with 4 mL PBS (Roche; 1666789) with 1% BSA (Sigma; A7888), neutralized with 65 µL 10N NaOH (Fisher; SS255-1), and vortexed for 30 seconds. The samples were then centrifuged at 4°C for 5 minutes at 2,000 RPM. The samples were diluted 5000X in PBS with 1% BSA, and insulin was measured (MSD; K152BZC-1).

#### RNA isolation and QC

RNA from both DO and CC-RIX adipose, gastrocnemius, and left liver lobe tissues was isolated using the MagMAX mirVana Total RNA Isolation Kit (ThermoFisher; A27828) and the KingFisher Flex purification system (ThermoFisher; 5400610). The frozen tissues were pulverized with a Bessman Tissue Pulverizer (Spectrum Chemical) and homogenized in TRIzol™ Reagent (ThermoFisher; 15596026) using a gentleMACS dissociator (Miltenyi Biotec Inc). After adding chloroform to the TRIzol homogenate, the RNA-containing aqueous layer was extracted for RNA isolation, following the manufacturer's protocol, starting with the RNA bead binding step using the RNeasy Mini kit (Qiagen; 74104). RNA concentrations and quality were assessed using the Nanodrop 8000 spectrophotometer (Thermo Scientific) and the RNA 6000 Pico or RNA ScreenTape assay (Agilent Technologies).

#### Library construction

Before library construction, 2 µL of diluted (1:1000) ERCC Spike-in Control Mix 1 (ThermoFisher; 4456740) was added to 100 ng of each RNA sample. Libraries were then constructed using the KAPA mRNA HyperPrep Kit (Roche Sequencing Store; KK8580) following the manufacturer's protocol. The process involves isolating polyA-containing mRNA using oligo-dT magnetic beads, fragmenting the RNA, synthesizing the first and second strands of cDNA, ligating Illumina-specific adapters with unique barcode sequences for each library, and performing PCR amplification. The quality and concentration of the libraries were evaluated using the D5000 ScreenTape (Agilent Technologies) and the Qubit dsDNA HS Assay (ThermoFisher; Q32851), respectively, according to the manufacturers' instructions.

#### Sequencing

Libraries were sequenced on an Illumina NovaSeq 6000 using the S4 Reagent Kit (Illumina; 20028312). All tissues underwent 100 bp paired-end sequencing, aiming for a target read depth of 30 million read pairs.

#### Trait selection in DO

We filtered the measured traits in this study to a set of relatively non-redundant measures that were well-represented in the population (having at least 80% of individuals measured). A complete description of trait filtering can be found at Figshare DOI: 10.6084/m9.figshare.27066979 in the file Documents > 1.DO > 1b.Trait_Selection.Rmd.

We took two approaches for traits with multiple redundant measurements, for example longitudinal body weights. In the case of longitudinal measurements, we used the final measurement, as this was the closest physiological measurement to the measurement of gene expression, which was done at the end of the experiment. The labels for these traits have the word "Final" appended to their name. For traits with multiple highly related measurements, such as cholesterol, we used the first principal component of the group of measurements. For example, we used the first principal component of all LDL measurements as the measurement of LDL. For each set of traits, we ensured the first principal component had the correct sign by correlating it with the average of the traits. For correlation coefficients (R) less than 0, we multiplied the principal component by -1. The labels for these traits have the term "PC1" appended to their name.

#### Processing of RNA sequencing data

We used the Expectation-Maximization algorithm for Allele Specific Expression (EMASE)[6;7] to quantify multi-parent allele-specific and total expression from RNA-seq data for each tissue. EMASE was performed by the Genotype by RNA-seq (GBRS) software package (https://gbrs.readthedocs.io/en/latest/). In the process, R1 and R2 FASTQ files were combined and aligned to a hybridized (8-way) transcriptome generated for the 8 DO founder strains as single-ended reads. GBRS was also used to reconstruct the mouse genotype probabilities along ~69K markers, which was used for confirming genotypes in the quality control (QC) process. For the QC process, we used a Euclidean distances method (developed by Greg Keele - Churchill Lab) to compare the GBRS genotype probabilities between the tissues and the genotype probabilities array for all mice. The counts matrix for each tissue was processed to filter out transcripts with less than one read for at least half of the samples. RNA-seq batch effects were removed by regressing out batch as a random effect and considering sex and generation as fixed effects using lme4 R package. RNA-Seq counts were normalized relative to total read counts using the variance stabilizing transform (VST) as implemented in DESeq2 and using rank normal score.

#### eQTL analysis

We used R/qtl2[8] to perform eQTL analysis. We used the rank normal score data and used sex and DO generation as additive covariates. We also used kinship as a random effect. We used permutations to find a LOD threshold of 8 for significant QTLs which corresponded to a $p$ value of 0.05.

To assess whether eQTL were shared across tissues, we considered significant eQTLs within 4Mb of each other to be overlapping. We considered local and distal eQTLs separately.

#### Local and distal heritability of transcripts

To estimate local and distal heritability of each transcript, we scaled each normalized transcript to have a variance of 1. We then modeled this transcript with the local genotype using the fit1() function in R/qtl with a kinship correction. We used the resulting model to predict the transcript values. The variance of the predicted transcript is its local heritability. We then estimated the heritability of the residual of the model fit. The variance of the residual multiplied by its heritability is the distal heritability of the transcript.

We compared local and distal estimates of heritability to measures of trait relevance for each transcript. Trait relevance was the Pearson correlation (R) between the transcript and the trait.

#### High-dimensional mediation analysis

In this section we derive the objective function for high-dimensional mediation analysis (HDMA) and present an iterative algorithm to optimize this objective function. Our starting point is the univariate case, where we describe perfect mediation as a constraint on the covariance matrix among variables. We then leverage this constraint to define projections of multivariate data that are maximally consistent with perfect mediation (HDMA). Next, we demonstrate how to *kernelize* HDMA to limit dimensionality of the model and enable non-linear HDMA models.

##### Perfect mediation as a constraint on covariance matrices

Suppose we have three random variables $x$, $m$, and $y$. Assume they each have unit variance and that they satisfy the following structural equation model (SEM) such that $m$ perfectly mediates the effect of $x$ on $y$:

$$m = \alpha x + \epsilon_m \tag{1}$$

$$y = \beta m + \epsilon_y \tag{2}$$

From these structural equations, we have the *model-implied covariance matrix*, $\Sigma$, given by

$$\Sigma = \begin{bmatrix} 1 & \alpha & \alpha\beta \\ \alpha & 1 & \beta \\ \alpha\beta & \beta & 1 \end{bmatrix} \tag{3}$$

Note that the assumption of perfect mediation forces the covariance between $x$ and $y$ to be $\alpha\beta$. In any finite data set, however, the observed covariance matrix, $S = [S_{ij}]$, will not typically satisfy this constraint.

The general negative log-likelihood fitting function for an SEM is given by

$$L = \mathrm{tr}\left(S\Sigma^{-1}\right) + \log|\Sigma|, \tag{4}$$

where $|\cdot|$ denotes the determinant of a matrix and $\mathrm{tr}(\cdot)$ denotes the trace[9]. For the perfect-mediation model, these values are

$$|\Sigma| = (1-\alpha^2)(1-\beta^2) \tag{5}$$

$$\Sigma^{-1} = \begin{bmatrix} 1/(1-\alpha^2) & -\alpha/(1-\alpha^2) & 0 \\ -\alpha/(1-\alpha^2) & (1-\alpha^2\beta^2)/\left((1-\alpha^2)(1-\beta^2)\right) & -\beta/(1-\beta^2) \\ 0 & -\beta/(1-\beta^2) & 1/(1-\beta^2) \end{bmatrix} \tag{6}$$

Plugging these into the likelihood function, we get

$$L = \log\left((1-\alpha^2)(1-\beta^2)\right) - \frac{2\alpha^2\beta^2}{(1-\alpha^2)(1-\beta^2)} + 1 - \frac{2\alpha}{1-\alpha^2}S_{12} - \frac{2\beta}{1-\beta^2}S_{23} \tag{7}$$

To simplify notation, we define

$$F(\alpha,\beta) = \log\left((1-\alpha^2)(1-\beta^2)\right) - \frac{2\alpha^2\beta^2}{(1-\alpha^2)(1-\beta^2)} + 1, \tag{8}$$

so the likelihood function is now

$$L = F(\alpha,\beta) - \frac{2\alpha}{1-\alpha^2}S_{12} - \frac{2\beta}{1-\beta^2}S_{23} \tag{9}$$

Note that this likelihood is maximized by fitting regression coefficients $\alpha$ and $\beta$ between $x$ and $m$ and $m$ and $y$, respectively, but the negative log-likelihood formulation is useful for the multivariate extension below.

**Projecting multivariate data to identify latent mediators**

Suppose now that we have three data matrices, $X$, $M$, and $Y$ (individuals by variables) that are mean centered by column. The central assumption of HDMA is that these multivariate data encode *latent variables* that are causally linked according to the perfect-mediation model, in a sense made precise as follows.

We use the log-likelihood function (Eqn. 7) of the perfect mediation model as an objective function to identify latent variables, $l_X$, $l_M$, and $l_Y$, that are are correlated as closely as possible to the constraints of the perfect mediation model, Eqn. (3). We estimate these latent variables as linear combinations of the measured variables

$$l_X = Xa \tag{10}$$

$$l_M = Mb \tag{11}$$

$$l_Y = Yc \tag{12}$$

The coefficient vectors $a$, $b$, and $c$, are called *loadings*, analogous to the terminology in PCA and CCA. Because the data matrices are mean centered, we have

$$\text{mean}(l_X) = \text{mean}(l_M) = \text{mean}(l_Y) = 0, \tag{13}$$

and we assume the loadings are scaled so that each latent variable has unit variance

$$\text{var}(l_X) = \text{var}(l_M) = \text{var}(l_Y) = 1. \tag{14}$$

Plugging these formulae into the objective function (Eqn. 9), we have

$$S_{12} = \text{corr}\,(l_X, l_M) \tag{15}$$

$$S_{23} = \text{corr}\,(l_M, l_Y) \tag{16}$$

$$L(\alpha, \beta, a, b, c) = F(\alpha, \beta) - \frac{2\alpha}{1-\alpha^2}\text{corr}\,(l_X, l_M) - \frac{2\beta}{1-\beta^2}\text{corr}\,(l_M, l_Y) \tag{17}$$

$$= F(\alpha, \beta) - \frac{2\alpha}{1-\alpha^2}\text{corr}\,(Xa, Mb) - \frac{2\beta}{1-\beta^2}\text{corr}\,(Mb, Yc) \tag{18}$$

This yields an objective function of two sets of parameters: the *structural parameters* $\alpha$ and $\beta$ that define the causal model among latent variables, and the loading vectors $a$, $b$, and $c$, that define the latent variables in terms of the measured variables. The goal of HDMA is to optimize $L$ as a function of all parameters simultaneously. The form of the objective function, Eqn. 17, is effectively a weighted sum of correlation coefficients, connecting it to so-called *sum-of-correlation*, or SUMCOR, optimization problems[10], which we discuss further below.

#### An algorithm for HDMA

The global optimization of 17 is challenging because it is not a convex problem. However, the decomposition of the variables into structural and loading variables suggests an iterative algorithm, similar to the expectation-maximization algorithm, that converges at least to a stationary point. The overall idea is to use a block-coordinate-ascent strategy that iterates between optimizing $a$, $b$, and $c$, then optimizing $\alpha$ and $\beta$.

For fixed $a$, $b$, and $c$, the optimal $\alpha$ and $\beta$ are simply given by regression coefficients between $l_X$ and $l_M$ and $l_M$ and $l_Y$, respectively. Given these regression coefficients, $\alpha$ and $\beta$, we then optimize $a$, $b$, and $c$. For fixed $\alpha$ and $\beta$, the term $F(\alpha, \beta)$ is irrelevant, so minimizing the negative log-likelihood function reduces to maximizing the reduced function

$$L_{red}(a, b, c) = \frac{2\alpha}{1-\alpha^2}\text{corr}\left(Xa, Mb\right) + \frac{2\beta}{1-\beta^2}\text{corr}\left(Mb, Yc\right), \tag{19}$$

which is a weighted sum of correlation coefficients. This is exactly a (weighted) SUMCOR optimization problem[10]. These optimization problems are still not convex, but Tenenhaus *et al.* have recently proved convergence for iterative algorithms that optimize weighted SUMCOR problems[10–12]. These algorithms only guarantee convergence to a stationary point not necessarily a maximum, as is common in other non-convex problems, but this can be overcome with multiple random restarts, if needed. Thus, we have a sub-routine `wSUMCOR`$(X, M, Y, w_1, w_2)$ that solves the weighted SUMCOR problem

$$L_{wSUMCOR}(a, b, c, w_1, w_2) = w_1\text{corr}\left(Xa, Mb\right) + w_2\text{corr}\left(Mb, Yc\right). \tag{20}$$

Iterating between optimizing the structural parameters and loading parameters, we reduce the negative log-likelihood at each step and converge to a fixed point.

We summarize our optimization procedure in Algorithm 1.

#### Kernel HDMA

For large data matrices $X$, $M$, and $Y$, especially with high correlation among variables, as is common for high-throughput biological assays (*e.g.*, ~1M alleles for genotypes, ~20k transcripts), we can further reduce the dimensionality of the HDMA model by requiring that loading vectors lie in the span of the the measured

**Algorithm 1** High-dimensional mediation analysis

**Input:** $X$, $M$, $Y$ ▷ Data matrices
**Output:** $\alpha$, $\beta$, $a$, $b$, $c$, $l_X$, $l_M$, $l_Y$ ▷ Structural parameters, loadings, scores
  $\alpha \leftarrow 0.5$, $\beta \leftarrow 0.5$ ▷ Initialize structural parameters
  **while** $converge \neq TRUE$ **do**
    $d \leftarrow \frac{2\alpha}{1-\alpha^2} + \frac{2\alpha}{1-\alpha^2}$ ▷ Normalization constant for weights
    $w_1 \leftarrow \frac{1}{d}\frac{2\alpha}{1-\alpha^2}$, $w_2 \leftarrow \frac{1}{d}\frac{2\beta}{1-\beta^2}$ ▷ Set weights (sum to one)
    $(a, b, c) \leftarrow \texttt{wSUMCOR}(X, M, Y, w_1, w_2)$ ▷ Compute loadings
    $l_X \leftarrow Xa$, $l_M \leftarrow Mb$, $l_Y \leftarrow Yc$ ▷ Compute scores
    $\alpha \leftarrow \mathrm{corr}(l_X, l_M)$, $\beta \leftarrow \mathrm{corr}(l_M, l_Y)$ ▷ Update structural parameters
  **end while**

individuals, namely

$$a = X^T\tilde{a} \tag{21}$$

$$b = M^T\tilde{b} \tag{22}$$

$$c = Y^T\tilde{c}. \tag{23}$$

This replaces the full feature data, say $X$, with the covariances among individuals (aka, Gram matrices), $C_X = XX^T$, and reduces the dimensionality from the number of measured variables down to the number of individuals

$$l_x = XX^T\tilde{a} = C_X\tilde{a} \tag{24}$$

$$l_M = MM^T\tilde{b} = C_M\tilde{b} \tag{25}$$

$$l_M = YY^T\tilde{c} = C_Y\tilde{c}. \tag{26}$$

This reduction is called *kernelization*[12] and is widely applied to other linear models, including CCA, linear regression, and classification.

It is interesting to note that kernelization is often used to convert a linear model to a non-linear model by replacing the covariance matrices, *e.g.* $C_X$, with more complex *kernel matrices* $K_X$ that encode similarity measures among individuals that are non-linear functions of the measured variables. non-linear model by replacing the covariance matrices, *e.g.* $C_X$, with more complex *kernel matrices* $K_X$ that encode similarity measures among individuals that are non-linear functions of the measured variables. Promoting a linear model to a non-linear model in this way is called the *kernel trick* and is widely used in the machine learning field. The above considerations show that HDMA is kernelizable in the same way as other linear models, although the exploration of non-linear models is outside the scope of this study.

##### Implementation details

We have implemented HDMA (Algorithm 1) in the R programming language. Tenenhaus *et al.* have implemented their optimizers in the Regularized Generalized Canonical Correlation Analysis (RGCCA) R package[13], which we use as the subroutine `wSUMCOR`. As Tenenhaus *et al.* discuss optimizing the empirical correlation coefficient *per se* is numerically unstable due to the inversion of the covariance matrices of the measured variables (*e.g.*, the transcript-transcript covariance matrix). To overcome this, the RGCCA package uses a regularized form of the covariance matrix developed by Schaeffer and Strimmer[14], which can be estimated rapidly using an analytic formula.

As a convergence criterion, we stop the iterations when both $\alpha$ and $\beta$ change by less than $10^{-6}$ from their previous value in one iteration.

All code required to run HDMA is available at Figshare: https://figshare.com/ DOI: 10.6084/m9.figshare.27066979

#### Enrichment of biological terms

We performed gene set enrichment analysis (GSEA)[15] using the transcript loadings in each tissue as gene weights. GSEA determines enrichment of pathways based on where the contained genes appear in a ranked list of genes. If the genes in the pathway are more concentrated near the top (or the bottom) of the list than expected by chance, the pathway can be interpreted as being enriched with positively (negatively) loaded transcripts. We used the R package fgsea[16] to calculate normalized enrichment scores for all GO terms and all KEGG pathways.

We downloaded all KEGG[17] pathways for *Mus musculus* using the R package clusterProfiler[17]. We then used fgsea to calculate enrichment scores in each tissue using the transcript loadings in each tissue as our ranked list of genes. We reported the normalized enrichment score (NES) for the 10 pathways with the largest positive NES and the 10 pathways with the largest negative NES.

We used the R package pathview[18] to visualize the loadings from each tissue in interesting pathways. We scaled the loadings in each tissue by the maximum absolute value of loadings across all tissues to compare them across tissues.

We downloaded GO term annotations from Mouse Genome Informatics at the Jackson Laboratory[19] https:

//www.informatics.jax.org/downloads/reports/index.html We removed gene-annotation pairs labeled with NOT, indicating that these genes were known not to be involved in these GO terms. We also limited our search to GO terms with between 80 and 3000 genes. We used the R package annotate[20] to identify the ontology of each term and the R package pRoloc[21] to convert between GO terms and names. As with the KEGG pathways, we used fgsea to calculate a normalized enrichment score for each GO term and collected loadings for the transcripts in each term to compare across tissues.

#### TWAS in DO mice

We performed a transcriptome-wide analysis (TWAS)[22;23] in the DO mice to compare to the results of high-dimensional mediation. To perform TWAS, we fit a linear model to explain variation in each transcript across the population using the genotype at the nearest marker to the gene transcription start site (TSS). We used kinship as a random effect and sex, diet, and DO generation as fixed effects. The predicted transcript from each of these models was the imputed transcript based only on the local genotype.

We correlated each imputed transcript with each of the metabolic phenotypes after adjusting phenotypes for sex, diet, and DO generation. To calculate significance of these correlations, we performed permutation testing by shuffling labels of individual mice and recalculating correlation values. Significant correlations were those more extreme than any of the permuted values, corresponding to an empirical $p$ value of 0. These are transcripts whose locally encoded expression level was significantly correlated with one of the metabolic traits. This suggests an association between the genetically encoded transcript level and the trait but does not identify a direction of causation.

#### Literature support for genes

To determine whether each gene among those with large loadings or large heritability had a supported connection to obesity or diabetes in the literature, we used the R package easyPubMed[24]. We searched for the terms (“diabetes” OR “obesity”) along with the tissue name (adipose, islet, liver, or muscle), and the gene name. We restricted the gene name to appear in the title or abstract as some short names appeared coincidentally in contact information. We checked each gene with apparent literature support by hand to verify that support, and we removed spurious associations. For example, FAU is used as an acronym for fatty acid uptake and CAD is used as an acronym for coronary artery disease. Both terms co-occur with the terms diabetes and obesity in a manner independent of the genes *Fau* and *Cad*. Other genes that co-occurred with diabetes and obesity, but not as a functional connection were similarly removed. For example, the gene *Rpl27* is used as a reference gene for quantification of the expression of other genes, and co-occurrence with diabetes and obesity is a coincidence. We counted the abstracts associated with diabetes or obesity and each gene name and determined that a gene had literature support when it had at least two abstracts linking it to the terms diabetes or obesity in the respective tissue.

#### Tissue-specific clusters

To compare the top loading genes across tissues, we selected genes with a loading at least 2.5 standard deviations from the mean across all tissues. We made a matrix consisting of the union of these sets populated with the tissue-specific loading for each gene. We used the pam() function in the R package cluster[25] to cluster the loading profiles around $k$ medoids. We tested $k = 2$ through 20 and used silhouette andlysis to compare the separation of the clusters. The best separation was achieved with $k = 12$ clusters. For each cluster we used the R package gprofiler2[26] to identify enriched GO terms and KEGG pathways for the genes in each cluster.

#### CC-RIX genotypes

We used the most recent common ancestor (MRCA) genotypes for the Collaborative Cross (CC) mice available on the University of North Carolina Computational Systems Biology website: http://www.csbio.unc.edu/CCstatus/CCGenomes/

To generate CC-RIX genotypes, we averaged the haplotype probabilities for the two parental strains at each locus.

#### Imputation of gene expression in CC-RIX

To impute gene expression in the CC-RIX, we performed the following steps for each transcript in each tissue (adipose, liver, and skeletal muscle):

1. Calculate diploid CC-RIX genotype for all CC-RIX individuals at the marker nearest the transcription start site of the transcript.
2. Multiply the genotype probabilities by the eQTL coefficients identified in the DO population.

To check the accuracy of the imputation, we correlated each imputed transcript with the measured transcript. The average Pearson correlation (r) was close to 0.5 for all three tissues (Supp. Fig. S7A), and as expected, the correlation between the imputed transcript and the measured transcript was highly positively dependent on the local eQTL LOD score of the transcript (Supp. Fig. S7B).

#### Prediction of CC-RIX traits

We used both measured expression and imputed expression combined with the results from HDM in the to predict phenotype in the CC-RIX. The traits measured in the DO and the CC-RIX were not identical, so we limited our prediction to body weight, which was measured in both populations, and was the largest contributor to the phenotype score in the DO.

For each CC-RIX individual, we multiplied the transcript abundances across the transcriptome by the loadings derived from the HDM in the DO population (Fig. 7A). This resulted in a vector with $n$ elements, where $n$ is the number of transcripts in the trancriptome. Each element was a weighted value that combined the relative abundance of the transcript with how that abundance affected the phenotype. We averaged the values in this vector to calculate an overall predicted phenotype score for the individual CC-RIX animal.

After calculating this predicted phenotype value across all CC-RIX animals, we correlated the predicted values from each tissue with measured body weight (Fig. 7B).

#### Cell type specificity

We investigated whether the loadings derived from HDM reflected tissue composition changes in the DO mice prone to obesity on the high-fat diet. To do this, we acquired lists of cell-type specific transcripts from the literature. In adipose tissue, we looked at cell-type specific transcripts for macrophages, leukocytes, adipocyte progenitors, and adipocytes as defined in [29087381]. In pancreatic islets, we looked at cell-type specific transcripts for alpha cells, beta cells, delta cells, ductal cells, mast cells, macrophages, acinar cells, stellate cells, gamma and epsilon cells, and endothelial cells as defined by [36778506]. Both studies defined cell-type specific transcripts based on human cell types. We collected the loadings for each set of cell-type specific transcripts in the respective tissue and asked whether the mean loading for the cell type differed significantly from 0 (Fig. 8). A significant positive loading for the cell type would suggest a genetic predisposition to have a higher proportion of that cell type in the tissue. To determine whether each mean loading differed significantly from 0, we performed permutation tests. We randomly sampled $n$ genes outside of the cell-type specific, where $n$ was the number of genes in the set. We compared the distribution of loading means over 10,000 random draws to that seen in the observed data. We used a significance threshold of 0.01.

#### Comparison of transcriptomic signatures to human transcriptomic signatures

To compare the transcriptomic signatures identified in the DO mice to those seen in human patients, we downloaded human gene expression data from the Gene Expression Omnibus (GEO)[27;28]. We focused on adipose tissue because this had the strongest relationship to obesity and insulin resistance in the DO. We downloaded the following human gene expression data sets:

- Accession number GSE152517 - Performed bulk RNA sequencing on visceral adipose tissue resected from seven diabetic and seven non-diabetic obese individuals.
- Accession number GSE44000 - Used Agilent-014850 4X44K human whole genome platform arrays (GPL6480) to measure gene expression in purified adipocytes derived from the subcutaneous adipose tissue of seven obese (BMI>30) and seven lean (BMI<25) post-menopausal women.
- Accession number GSE205668 - Subcutaneous adipose tissue was resected during elective surgery from 35 normal weight, and 26 obese children. Gene expression was measured by RNA sequencing with an Illumina HiSeq 2500.
- Accession number GSE29231 - Visceral adipose biopsies were taken from three female patients with type 2 diabetes, and three non-diabetic female patients. Expression was measured with Illumina HumanHT-12 v3 Expression BeadChip arrays.

We downloaded each data set from GEO using the R package GEOquery[29]. In each case, we verified that gene expression was log transformed and performed the transformation ourselves if it had not already been done. When covariates such as age and sex were available in the metadata files, we regressed out these variables. We mean centered and standardized gene expression across transcripts.

We matched the human gene expression to the mouse gene expression by pairing orthologs as defined in The Jackson Laboratory's mouse genome informatics data base (MGI)[30]. We multiplied each transcript in the human data by the adipose tissue loading of its ortholog in the DO mice. This resulted in a vector of weighted transcript values for each patient based on their own transcriptional profile and the obesity-related transcriptional signature from the DO analysis. The mean of this vector for an individual was the prediction of their obesity status. Higher values indicate a prediction of higher obesity or risk of metabolic disease based on adipose gene expression. We then compared the values across groups, either obese and non-obese, or diabetic and non-diabetic depending on the groups in each study.

#### Connectivity Map Queries

We queried the transcript loading signatures from adipose tissue and pancreatic islets with the CMAP database. These tissues are the most related to metabolic disease and diabetes respectively.

The gene expression profiles in the Connectivity Map database are derived from human cell lines and human primary cultures and are indexed by Entrez gene IDs. To query the CMAP database, we identified the Entrez gene IDs for the human orthologs of the mouse genes expressed in each tissue. Each CMAP query takes the 150 most up-regulated and the 150 most down-regulated genes in a signature, however, not all human genes are included in their database. To ensure we had as many genes as possible in the query, we selected the top and bottom 200 genes with the most extreme positive and negative loadings respectively. We pasted these into the CLUE query application available at https://clue.io/query.

We filtered the results in two ways: First, we looked at the most significantly anti-correlated ($-log_{10}(\mathrm{FDR}q) > 15$) hits across all cell types. Second, we looked at the most anti-correlated within the most related cell type to the query and considered hits regardless of $-log_{10}(\mathrm{FDR}p)$. For adipose tissue we looked in normal adipocytes, abbreviated ASC in the CMAP database, and for pancreatic islets we looked in pancreatic cancer cells, abbreviated YAPC in the CMAP database.

[24] Damiano Fantini. *easyPubMed: Search and Retrieve Scientific Publication Records from PubMed*, 2019. R package version 2.13.

[25] Martin Maechler, Peter Rousseeuw, Anja Struyf, Mia Hubert, and Kurt Hornik. *cluster: Cluster Analysis Basics and Extensions*, 2023. R package version 2.1.6 — For new features, see the 'NEWS' and the 'Changelog' file in the package source).

[26] Liis Kolberg, Uku Raudvere, Ivan Kuzmin, Jaak Vilo, and Hedi Peterson. gprofiler2– an r package for

gene list functional enrichment analysis and namespace conversion toolset g:profiler. *F1000Research*, 9 (ELIXIR)(709), 2020. R package version 0.2.3.

[27] E. Clough, T. Barrett, S. E. Wilhite, P. Ledoux, C. Evangelista, I. F. Kim, M. Tomashevsky, K. A. Marshall, K. H. Phillippy, P. M. Sherman, H. Lee, N. Zhang, N. Serova, L. Wagner, V. Zalunin, A. Kochergin, and A. Soboleva. NCBI GEO: archive for gene expression and epigenomics data sets: 23-year update. *Nucleic Acids Res*, 52(D1):D138–D144, Jan 2024.

[28] R. Edgar, M. Domrachev, and A. E. Lash. Gene Expression Omnibus: NCBI gene expression and hybridization array data repository. *Nucleic Acids Res*, 30(1):207–210, Jan 2002.

[29] Sean Davis and Paul Meltzer. Geoquery: a bridge between the gene expression omnibus (geo) and bioconductor. *Bioinformatics*, 14:1846–1847, 2007.

[30] R. M. Baldarelli, C. L. Smith, M. Ringwald, J. E. Richardson, C. J. Bult, A. Anagnostopoulos, D. A. Begley, S. M. Bello, K. Christie, J. H. Finger, P. Hale, T. F. Hayamizu, D. P. Hill, M. N. Knowlton, D. M. Krupke, M. McAndrews, M. Law, I. J. McCright, L. Ni, H. Onda, D. Sitnikov, C. M. Smith, M. Tomczuk, L. Wilming, J. Xu, Y. Zhu, O. Blodgett, J. W. Campbell, L. E. Corbani, P. Frost, S. C. Giannatto, D. B. Miers, H. Motenko, S. B. Neuhauser, D. R. Shaw, N. E. Butler, and J. E. Ormsby. Mouse Genome Informatics: an integrated knowledgebase system for the laboratory mouse. *Genetics*, 227(1), May 2024.
